# AlphaFold2 models of the active form of all 437 catalytically competent human protein kinase domains

**DOI:** 10.1101/2023.07.21.550125

**Authors:** Bulat Faezov, Roland L. Dunbrack

## Abstract

Humans have 437 catalytically competent protein kinase domains with the typical kinase fold, similar to the structure of Protein Kinase A (PKA). Only 155 of these kinases are in the Protein Data Bank in their active form. The active form of a kinase must satisfy requirements for binding ATP, magnesium, and substrate. From structural bioinformatics analysis of 40 unique substrate-bound kinases, we derived several criteria for the active form of protein kinases. We include requirements on the DFG motif of the activation loop but also on the positions of the N-terminal and C-terminal segments of the activation loop that must be placed appropriately to bind substrate. Because the active form of catalytic kinases is needed for understanding substrate specificity and the effects of mutations on catalytic activity in cancer and other diseases, we used AlphaFold2 to produce models of all 437 human protein kinases in the active form. This was accomplished with templates in the active form from the PDB and shallow multiple sequence alignments of orthologs and close homologs of the query protein. We selected models for each kinase based on the pLDDT scores of the activation loop residues, demonstrating that the highest scoring models have the lowest or close to the lowest RMSD to 22 non-redundant substrate-bound structures in the PDB. A larger benchmark of all 130 active kinase structures with complete activation loops in the PDB shows that 80% of the highest-scoring AlphaFold2 models have RMSD < 1.0 Å and 90% have RMSD < 2.0 Å over the activation loop backbone atoms. Models for all 437 catalytic kinases are available at http://dunbrack.fccc.edu/kincore/activemodels. We believe they may be useful for interpreting mutations leading to constitutive catalytic activity in cancer as well as for templates for modeling substrate and inhibitor binding for molecules which bind to the active state.

## INTRODUCTION

Protein kinases regulate most cellular processes in eukaryotes. In humans, their dysregulation is often involved in disease and they are therefore often targets in drug development, especially in cancer (Cohen, Cross et al. 2021). A large majority of human protein kinases take on a common fold first determined by Susan Taylor and colleagues in 1991 (Knighton, Zheng et al. 1991), consisting of an N-terminal domain of five beta strands and the C-helix, and a largely helical C-terminal domain. The residues involved in catalytic activity are contained in the catalytic and activation loops that form a pocket for ATP binding and a groove for substrate binding in between the N and C terminal domains. Humans have 481 genes which contain at least one typical full-length protein kinase domain; 13 of these have two kinase domains, for a total of 494 kinase domains (Modi and Dunbrack 2019) [NB: since that paper was published, three kinases have been determined to be pseudogenes]. Of these, 437 are likely catalytic kinases, participating in phosphorylation of Ser, Thr, or Tyr residues on proteins, and 57 are likely pseudokinases. Currently in the PDB there are structures for 292 human typical kinase domains, of which 268 are catalytic kinases and 24 are pseudokinases (Modi and Dunbrack 2022).

Active and inactive conformations of typical kinases have been classified in several ways (Jacobs, Caron et al. 2008, Hari, Merritt et al. 2013, Ung, Rahman et al. 2018, Modi and Dunbrack 2019, Kanev, de Graaf et al. 2021). The active form is generally very similar across kinases because of the requirements of binding ATP, magnesium ions, and substrate, which impart constraints on the conformation of the activation loop (Johnson, Noble et al. 1996). Early on in the history of structure determination of kinases, a classification of structures into “DFGin” and “DFGout” was described (Levinson, Kuchment et al. 2006). In DFGin structures, the Asp side chain of the DFG motif is “in” the ATP binding site and the Phe side chain of the DFG motif is in a pocket under or adjacent to the C-helix of the N-terminal domain. In DFGout structures, the Asp side chain is “out” of the active site and the Phe side chain is removed from the C-helix pocket, allowing for the binding of Type 2 inhibitors such as imatinib that span both the ATP site and the C-helix pocket (Schindler, Bornmann et al. 2000).

There are, however, additional requirements for substrate binding and kinase activity (Johnson, Noble et al. 1996, Lowe, Noble et al. 1997). We previously used the presence of bound ATP, magnesium ion, and a phosphorylated activation loop to identify a set of 24 “catalytically primed” structures of 12 different kinases in the PDB (Modi and Dunbrack 2019). We found that in addition to being “DFGin,” these structures possess specific backbone and side-chain dihedral angles for the DFG motif (“*BLAminus*”), including the backbone dihedral angles of the residue immediately preceding DFG and the side-chain ξ1 dihedral angle of the DFG-Phe residue. They also possess a well-characterized salt bridge between a conserved glutamic acid residue in the C-helix and a conserved lysine residue in beta strand 3 of the N-terminal domain (Yang, Wu et al. 2012). These structures are often referred to as “C-helix-in.” Using these criteria for active kinases, only 183 of 437 catalytic typical human kinase domains are currently represented in the PDB with active structures. Additional criteria on the positions of the N-terminal and C-terminal segments of the activation loop (see Results) reduce this number to 155 kinases or 35%. Only 130 of 437 catalytic human kinases (30%) possess active structures and complete coordinates for the activation loop.

The program AlphaFold2 from DeepMind is a deep-learning program for highly accurate protein structure prediction and is trained on a large number of structures from the PDB (Jumper, Evans et al. 2021). It uses as input the query sequence, a multiple sequence alignment (MSA) of homologs of the query, and optionally template structures related to the query. DeepMind has provided models of nearly all human proteins produced by AlphaFold2, which are available on a website provided by the European Bioinformatics Institute (Varadi, Anyango et al. 2022). However, only 209 of the 437 (48%) catalytic human protein kinases have a fully active model in the EBI data set.

Because of the importance of knowing the active-state structures of kinases for understanding such features as substrate recognition, the effect of activating mutations in cancer, and drug development, in this paper we describe a pipeline for producing active models of typical protein kinases using the program AlphaFold2. Several groups have found that using MSAs of reduced depth and templates in specific conformational states coerces AF2 to produce conformationally variable models, including some models in the conformational state of the templates (Del Alamo, Sala et al. 2022, Heo and Feig 2022). We use similar techniques to compute predicted structures of active kinases.

A key aspect of this work is that we utilize structural bioinformatics of 40 non-redundant substrate-kinase complexes from the PDB to define strict criteria for identifying catalytically active protein kinases structures, including both experimental structures and models predicted by AlphaFold2. We impose criteria on the position of the Phe residue and the dihedral angles of the DFG motif, the formation of the N-terminal domain salt bridge (in kinases that possess the appropriate residues), and on the positions of the N and C terminal halves of the activation loop necessary for the formation of a substrate binding cleft. In addition to reduced MSAs from various sources and active templates from the PDB, we use catalytically active models of kinases produced by AF2 as additional templates for kinases which are more recalcitrant in producing active models and for additional sampling for all kinases. We refer to these as “distillation templates” in analogy with predicted structures that AF2 was trained on (“the distillation training set” (Jumper, Evans et al. 2021)).

We benchmark our protocol with 22 substrate-bound kinase structures in the PDB with complete activation loops and a set of 130 kinase structures from the PDB with complete activation loops that satisfy our active criteria. We show that the pLDDT scores for the activation loop are inversely correlated with RMSD of the activation loop for well characterized kinases. With these methods, we have produced active models of all 437 catalytic human protein kinase domains and made these models available on our Kinase Conformational Resource website and database, KinCore (http://dunbrack.fccc.edu/kincore/activemodels).

## RESULTS

### Catalytic protein kinases

To make catalytically active models of all human kinases with typical protein kinase domains, we need to distinguish between catalytic protein kinase domains and non-catalytic protein kinase domains or pseudokinases. *Catalytic* protein kinase domains are those able to phosphorylate proteins on Ser, Thr, or Tyr residues. Non-catalytic protein kinase domains or pseudokinases are domains that possess the typical protein kinase fold but lack protein kinase activity, although they may have other catalytic activity (e.g., PAN3, POMK). We previously published an alignment of all 497 human kinase domains from 484 genes annotated by Uniprot (Modi and Dunbrack 2019). This list excludes atypical kinases, such as ADCK, PI3/PI4, Alpha, FAST, and RIO kinases (https://www.uniprot.org/docs/pkinfam.txt). Since that time, three kinase genes have been identified as likely pseudogenes (SIK1B, PDPK2P, and PRKY) (Frankish, Carbonell-Sala et al. 2023), leaving us with 481 genes and 494 domains.

We identified catalytic protein kinases based on the presence of the Asp residue in the HRD motif, the Asp residue in the DFG motif, and the Lysine residue of the N-terminal domain salt bridge. We reclassified WNK (“With No Lysine”) kinases as catalytic protein kinases. Several kinases were reclassified based on literature annotations (e.g., BUB1B/BUB1R is a pseudokinase (Suijkerbuijk, van Dam et al. 2012); RYK is a pseudokinase (Katso, Russell et al. 1999)). The result of these efforts was a list of 437 active kinase domains in 429 genes. Eight genes have two (likely) catalytic protein kinase domains: RPS6KA1, RPS6KA2, RPS6KA3, RPS6KA4, RPS6KA5, RPS6KA6, OBSCN, and SPEG. Our previous phylogenetic analysis classified these 437 active domains into families as follows: AGC (60 kinases), CAMK (83), CK1 (11), CMGC (65), NEK (11), OTHER (43), STE (45), TKL (37), and TYR (82).

On our Kincore website (http://dunbrack.fccc.edu/kincore) and in the text that follows, we use the family name as a prefix in front of the HUGO gene name (e.g., TYR_EGFR) (Seal, Braschi et al. 2023). The catalytic protein kinase domains and associated data are listed in Supplementary Table 1. The pseudokinase domains are listed in Supplementary Table 2.

### The characteristics of active protein kinase domains

To identify structural features of the active form of catalytic protein kinases, we created two sets of structures that constitute likely catalytically active structures of protein kinases. The first consists of structures in the Protein Data Bank of kinases with peptide or protein substrates bound at the active site (**Table 1**). The second consists of 391 structures of catalytic protein kinase structures (comprising 74 different kinases) with bound ATP, ADP, or an ATP analog that are also in the DFGin “*BLAminus*” conformational state of the DFGmotif that we found characteristic of “catalytically primed” kinase structures (Modi and Dunbrack 2019). The example shown in **Figure 1** is an active form of human AKT1 bound to a substrate, PDB:4ekk (Lin, Lin et al. 2012). To determine what features are important to catalytic activity, we compared the structures in these data sets to all available structures of kinases in the PDB (without ATP and/or not in the DFGin-*BLAminus* state).

**Figure 1.**
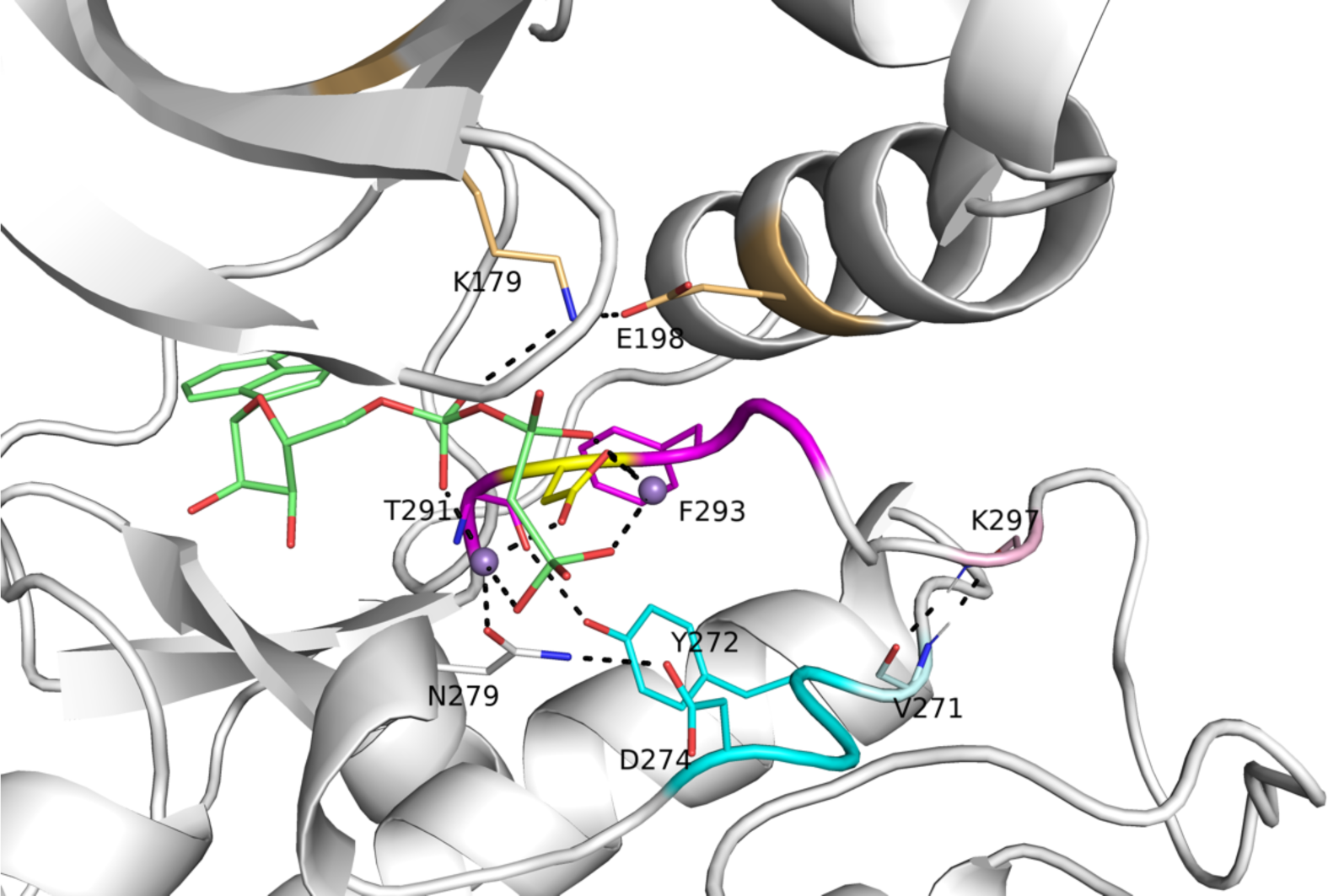
Active site of human AKT1 (PDB:4ekk, chain. **A).** Residues making hydrogen bonding interactions with ATP (green sticks), Mg^2+^ (purple spheres), the catalytic aspartic acid residue of the HRD motif (in AKT1, this is YRD; residues 272-274, cyan), and the aspartic residue (yellow) of the XDFG motif (residues 291-294, magenta) are shown in dashed lines. These include the salt bridge residues of the N-terminal domain (K179, E198, gold). Residue K297, which is the sixth residue of the activation loop (light pink), makes backbone-backbone hydrogen bonds with V271, which immediately precedes the YRD motif. In the stick representations, oxygen atoms are in red and nitrogen atoms are in blue. A substrate peptide is present in this structure but not shown in this figure.

**Table 1.**
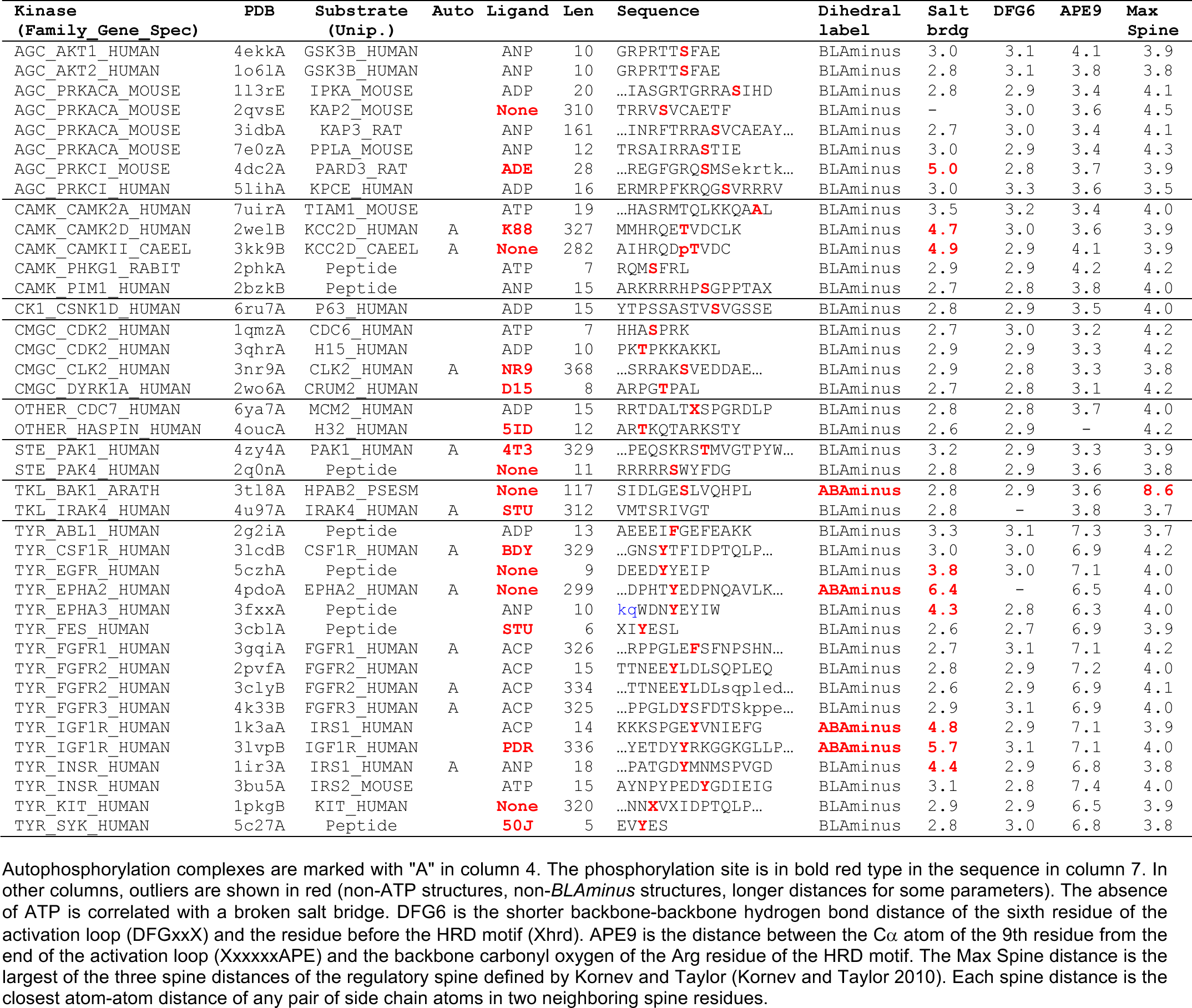
Kinase-substrate complexes in the Protein Data Bank (PDB)

Table 1 presents a list of unique kinase-substrate and kinase-pseudosubstrate complexes in the PDB and some structural parameters that will be considered below. Some of the “substrates” are in fact substrate-mimicking inhibitors, which bind similarly to substrates. Some kinases are represented more than once if they contain different bound substrates in the active site. Eleven of the 40 complexes are “autophosphorylation complexes,” which we previously identified as homodimeric complexes in crystals of protein kinases in which a known autophosphorylation site of one monomer sits in the active-site and substrate-binding groove of another monomer in the crystal (Xu, Malecka et al. 2015). These include autophosphorylation complexes of sites in the activation loop (STE_PAK1, 4zy4; TYR_IGF1R, 3lvp) and the kinase insert loop (TYR_FGFR1, 3gqi; TYR_FGFR3, 4k33). The remainder are N or C terminal tails (CAMK_CAMK2D, 2wel; CAMK_CAMKII, 3kk9; CMGC_CLK2, 3nr9, TYR_CSF1R, 3lcd; TYR_KIT, 1pkg; TYR_EPHA2, 4pdo; TYR_FGFR2, 3cly). Three other complexes are with larger proteins which are either direct substrates or inhibitors or both (AGC_PRKACA:KAP2, 2qvs; AGC_PRKACA:KAP3, 3idb; TKL_BAK1:HPAB2, 3tl8). The last of these is a plant kinase/pathogen-inhibitor complex (Cheng, Munkvold et al. 2011). The autophosphorylation complexes (marked with “A” in column 4 of Table 3) and inhibitor protein complexes provide insights of how kinases phosphorylate amino acids in the context of folded protein domains, as opposed to intrinsically disordered regions (IDRs) (Xu, Malecka et al. 2015).

We previously identified several criteria for active structures in the PDB for catalytic protein kinase domains (Modi and Dunbrack 2019): 1) the spatial label must be *DFGin*; 2) the dihedral label must be *BLAminus*; this indicates that the X, D, and F residues of the XDFG motif are in the “B”, “L”, and “A” regions of the Ramachandran map respectively, and the ξ1 rotamer of the Phe side chain is *g^-^* (∼ -60°); 3) there must be a salt bridge between the C-helix glutamic acid side chain and the beta strand 3 lysine side chain (the WNK kinases are an exception to this rule). In this paper, we validate these criteria and extend them to include: 4) the activation loop must be “extended” as determined by the presence of a backbone-backbone hydrogen bond between the sixth residue of the activation loop (X in DFGxxX) and the residue before the HRD motif (X in XHRD); 5) the C-terminal segment of the activation loop, which must be positioned for binding a substrate, as determined by a residue 9 positions from the end of the activation loop. We also consider the presence of the regulatory spine defined by Kornev and Taylor (Kornev and Taylor 2010). We review each of these in turn.

### DFGin conformation

The position of the DFG-Phe residue determines, in part, the position of the catalytic DFG-Asp residue. We defined DFGin by the distance between the DFG Phe Cζ atom and the Cα atoms of two residues in the N-terminal domain (Modi and Dunbrack 2019): the Lys residue in the β3 strand of the N-terminal domain salt bridge and the “Glu4” residue in the C-helix (Figure 1), which is the residue four residues following the Glu residue of the salt bridge. Based on these distances, structures are labeled as follows: *DFGin*, where the DFG-Phe residue is near the C-helix Glu4 residue but far from the Lys residue; *DFGout*, where the Phe residue is far from the C-helix Glu4 residue and close to the Lys residue; and *DFGinter*, where the Phe residue is not far from either the Glu4 or Lys residues. These distances are plotted for ATP and non-ATP-bound structures in **Figure 2**. The vast majority of ATP-bound structures (defined as having ligands with PDB 3-letter codes: ATP, ADP, ANP, or ACP in the active site) are *DFGin* with LysCα-PheCζ distance > 11 Å and Glu4Cα-PheCζ distance < 11 Å. All of the substrate-bound structures listed in Table 1 are DFGin (required for the *BLAminus* and *ABAminus* conformations of the XDF motif).

**Figure 2.**
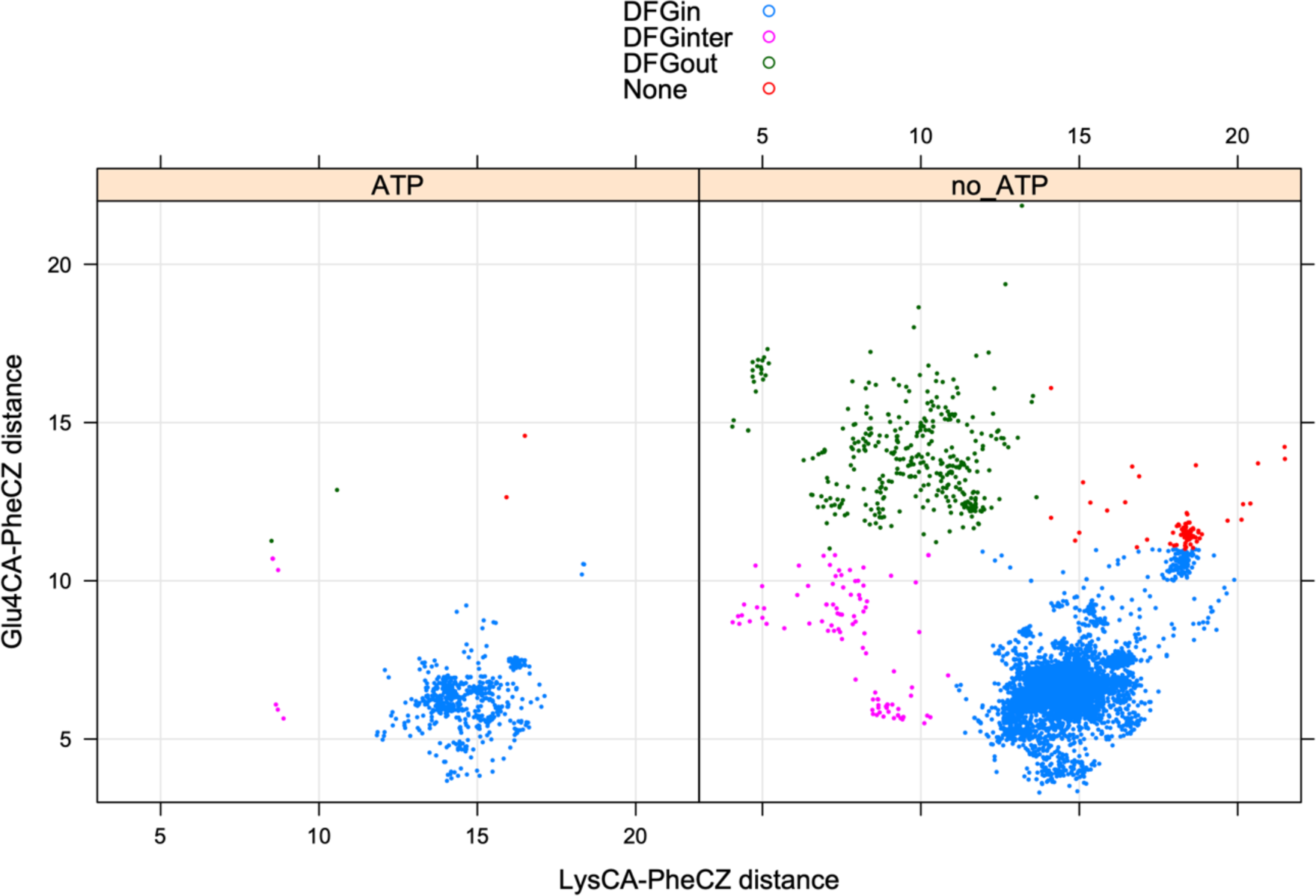
Defining DFGin and DFGout structures. Distances of the Cζ atom of DFG-Phe from the Cα atoms of the salt bridge Lys residue and Glu4 (4 residues after the salt bridge Glu).

### BLAminus conformation and Salt bridge formation

The conformation of the XDFG motif and the formation of the salt bridge in the N-terminal domain work together to form an active site capable of binding ATP and magnesium ions for the phosphorylation reaction. These interactions are shown in Figure 1, where the Asp of the DFG motif interacts with the active site magnesium ions which chelate ATP. The carbonyl oxygen of the residue before the DFG motif (X of XDFG, T291) forms a hydrogen bond with the Tyr residue of the YRD motif (usually HRD, but Tyr in AKT1). This hydrogen bond helps position the catalytic aspartic acid residue of the Y/HRD motif, which interacts with the Ser or Thr hydroxyl atoms of substrate residues to be phosphorylated. The *BLAminus* conformation is required for these interactions (Modi and Dunbrack 2019). *ABAminus* structures involve a “peptide flip” of the X-D residues (Hayward 2001), such that the carbonyl of the X residue points upwards and does not interact with Y/H of the Y/HRD motif. Many of these structures are missolved and should be *BLAminus*, as demonstrated by poor electron density for the X residue carbonyl oxygen (Modi and Dunbrack 2019). The backbone and side-chain dihedral angles of the XDFG motif for the substrate-bound structures in Table 1 are shown in Figure 3.

**Figure 3.**
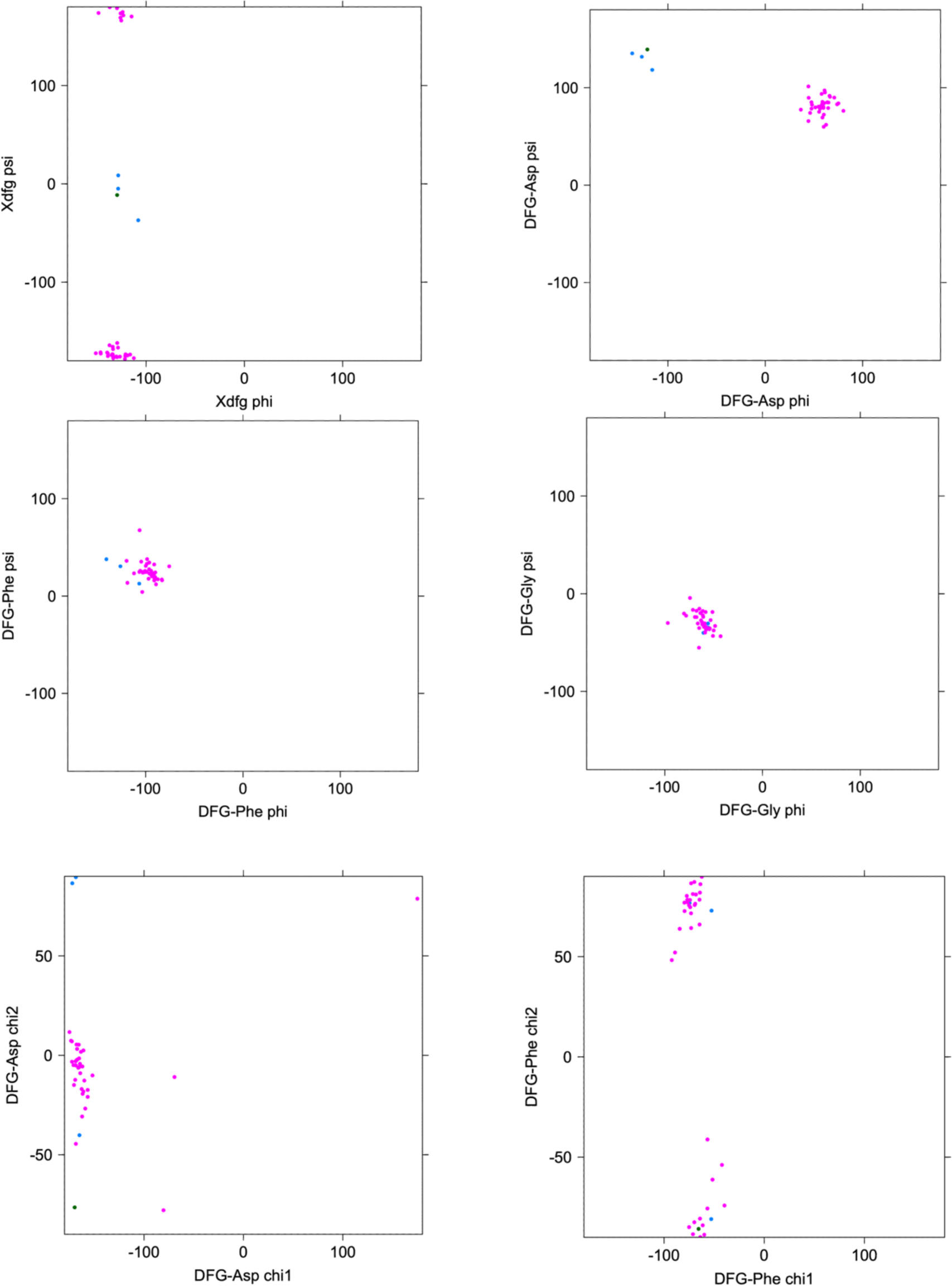
Ramachandran plots and side-chain dihedral angle plots for the XDFG motif residues in substrate-bound structures in Table 1. BLAminus structures are shown in magenta and ABAminus structures are shown in blue.

In addition, the Lys of the N-terminal domain salt bridge interacts directly with the alpha-beta phosphate linkage of ATP. The Glu of the salt bridge helps position the Lys in this interaction (Yang, Wu et al. 2012). The minus rotamer of the Phe side chain is required for this interaction, since the plus rotamer of the inactive *BLAplus* and *BLBplus* conformations points upwards (instead of downwards as in the *BLAminus* and *BLBminus* conformations) and pushes the C-helix outwards, breaking the salt bridge (Modi and Dunbrack 2019).

While 69% of ATP-bound and 65% of non-ATP-bound catalytic kinase structures are in the *BLAminus* conformation, the role of the *BLAminus* configuration becomes clearer when combining it with the formation of the N-terminal domain salt bridge. In **Figure 4A**, the density of distances of the salt bridge atom pairs (Nζ in the β3 Lys residue with Oχ1 or Oχ2 in the C-helix Glu residue (whichever is shorter)) is plotted for *BLAminus* and non-*BLAminus* structures with and without ATP. When *BLAminus* structures are bound with ATP, the salt bridge is strongly favored with a mean distance of about 3.0 Å (upper left of Figure 4A). However, ATP-bound structures that are not in the *BLAminus* state have a broken salt bridge, with most structures having a Lys/Glu distance greater than 10 Å (lower left panel of Figure 4A). Even in the absence of ATP, the *BLAminus* conformation encourages the formation of the salt bridge (upper right vs lower right panels of Figure 4A).

**Figure 4.**
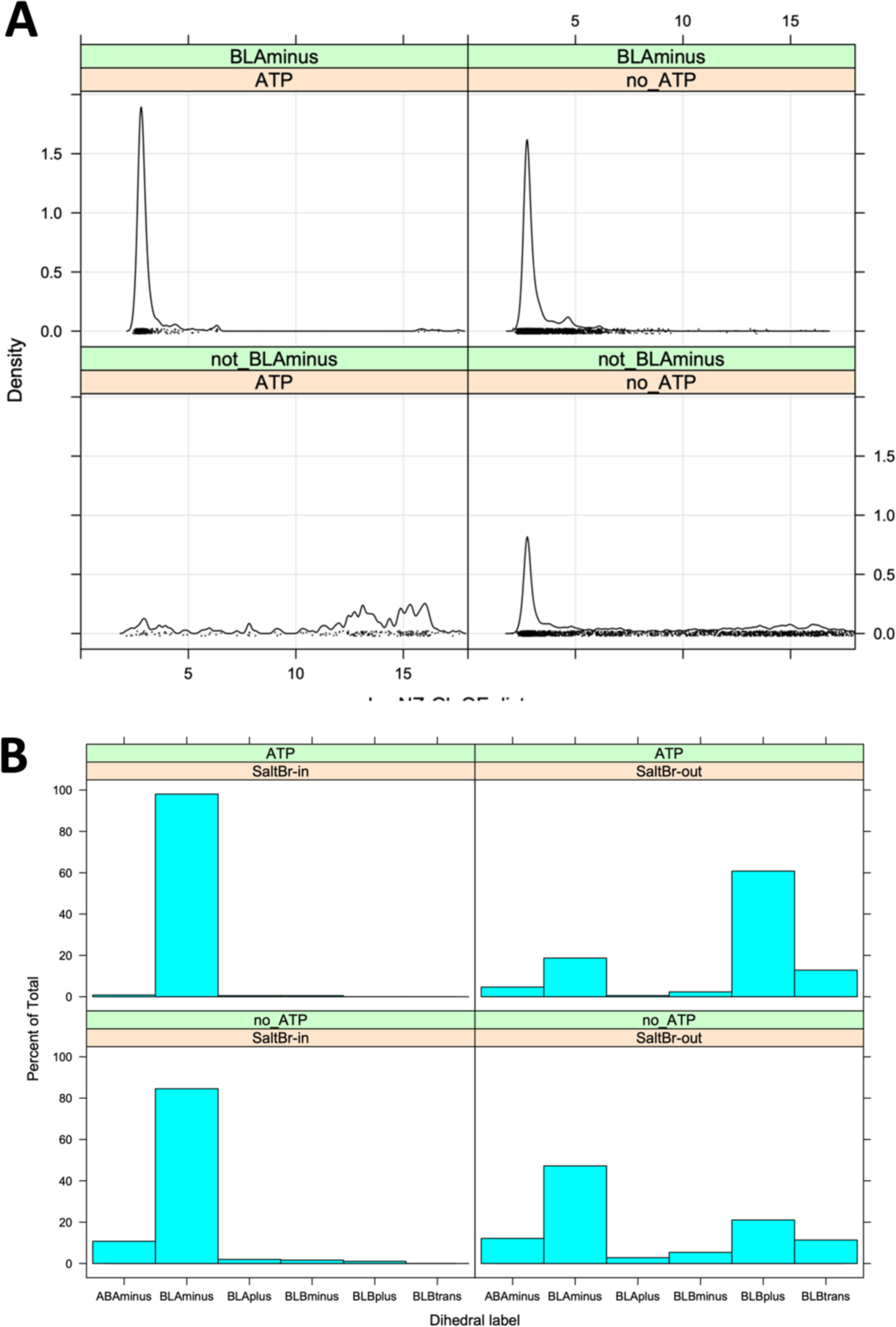
Relationship of the XDFG motif dihedral angle conformational state and the N-termainal domain salt bridge. **A.** Minimum distance of the Nζ atom from the Lys residue and the Oε1 or Oε2 atoms of the Glu residue of the salt bridge in structures with or without ATP and in or out of the BLAminus conformation. **B.** Distribution of dihedral angle states in ATP-bound and ATP-unbound structures in the presence of the absence of the N-terminal domain salt bridge (minimum Nζ/Oε distance cutoff 3.6 Å). When the salt bridge and ATP are present, 99% of structures are in the BLAminus conformation for the XDF motif.

Conversely, if we require salt bridge formation (“SaltBr-In”) with a cutoff of Nζ/Oε distance of 3.6 Å, 99% of ATP-bound structures are in the *BLAminus* conformation. When the salt bridge is not formed (“SaltBr-Out”), only 19% of the structures are *BLAminus* (**Figure 4B**).

### ActLoopNT

We examined the substrate-bound structures listed in Table 1 for further characteristics of the activation loop structure that may be required for binding substrates by determining contacts of the substrate with residues in the activation loop. These residues must be in the appropriate position for forming a substrate binding groove. Examples from four families are shown in **Figure 5** with the activation loops in magenta, phosphorylated residues in the activation loop in pink, ATP (or analogs) in green sticks, and the substrates in blue.

**Figure 5.**
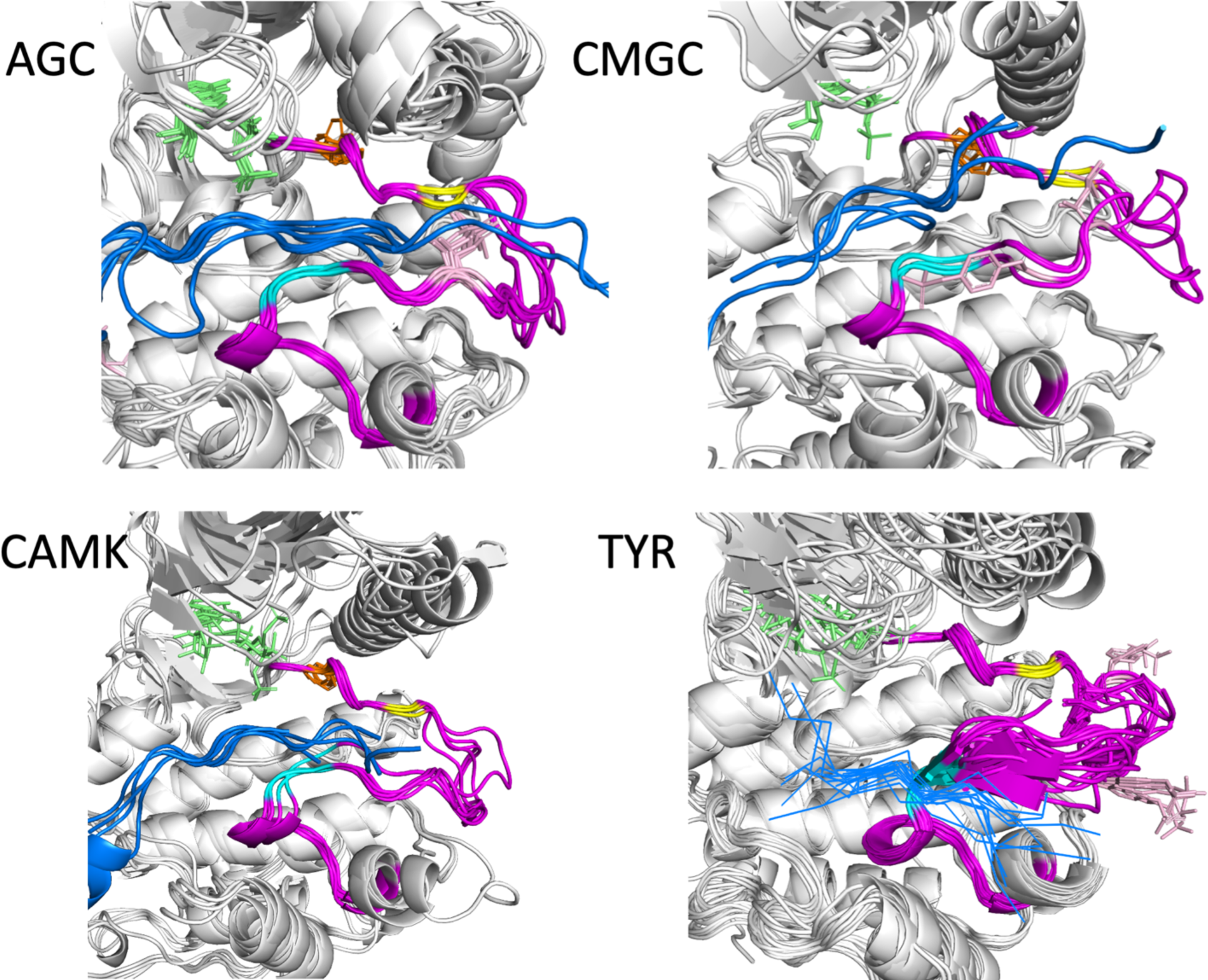
Substrate-bound structures in the AGC, CAMK, CMGC, and TYR families from Table 1. In each figure, the substrate peptides (or pieces of longer proteins) are in blue and the activation loop is in magenta. ATP or any analogue is shown in green sticks. The Phe of the DFG motif is shown in orange sticks and phosphorylated residues in the activation loop are in pink. The sixth residue of the activation loop (DFGxxX) is in yellow, while the 8^th^ and 9^th^ residues from the end of the activation loop are in cyan (XXxxxxAPE).

Besides the conformation of the DFG motif (the Phe/Tyr residue of DFG is shown in orange sticks), two other features are evident in the substrate-bound structures. The first is that the first few residues of the activation loop, up to at least the sixth residue (yellow in each figure), have similar conformations and positions across the members of each family. The second is that the C-terminal segment of the activation loop, up to at least 9 residues from the end of the activation loop, also shares a common conformation and position across family members. This segment is referred to as the “P+1 loop,” since it binds the side chain of the substrate residue immediately after the phosphorylation site (Lowe, Noble et al. 1997, Kornev and Taylor 2010). In Figure 5, residues 8 and 9 from the end of the activation loop are shown in cyan. In Ser/Thr kinases, the conformation of residues 8-11 from the end of the activation loop resemble the hull shape of an upside-down, round-bottom boat. Residues 8-9 in TYR kinases are also in a common position, although the structure diverges in residues 10 and 11 more than in the Ser/Thr kinase members. In TYR kinases, the substrate binds directly to these residues in the form of a short beta sheet (blue lines in Figure 5, lower right). The conformation in TYR kinases may diverge to accommodate substrates with larger or smaller side chains. In other kinase families, the substrate binds to a groove between the N and C-terminal segments of the activation loop.

To investigate potential requirements for substrate binding, we determined which residues within the activation loop form direct contacts with substrate residues (any atom contact within 5 Å between substrate residues and the DFG…APE sequence). The results are shown in **Table 2**. Most substrates have a contact with one or more of the DFG residues as well as the fourth residue of the activation loop, while a small number have contacts with residues 5 and 6. By looking at the structures, we identified the existence of backbone-backbone hydrogen bonds between residue 6 of the activation loop (DFGxx***X***) and the residue immediately preceding the HRD motif (***X***HRD) (**Figure 6A**). This backbone-backbone hydrogen bond is contained in two very short anti-parallel beta strands (3 residues each), labeled beta strand 6 (comprising the three residues preceding the HRD motif) and beta strand 9 (comprising residues 6-8 of the activation loop) in protein kinase structures (Kornev and Taylor 2010). We used this hydrogen bond previously to characterize active structures in the PDB (Modi and Dunbrack 2019). This hydrogen bond is present in all of the substrate-bound structures in Table 1 (“DFG6” in the table) except for IRAK4 (PDB:4u97) where the DFG6 residue is disordered. Almost 99% of *BLAminus* structures (386 of 391) with bound ATP contain this hydrogen bond with the minimum N-O or O-N backbone-backbone distance less than 3.6 Å (**Figure 6B**, upper left panel). The only exception is an activation-loop swapped structure of STE_MAP4K1 (PDB: 6cqd) in which the N-terminal portion of the activation loop forms an α-helix.

**Figure 6.**
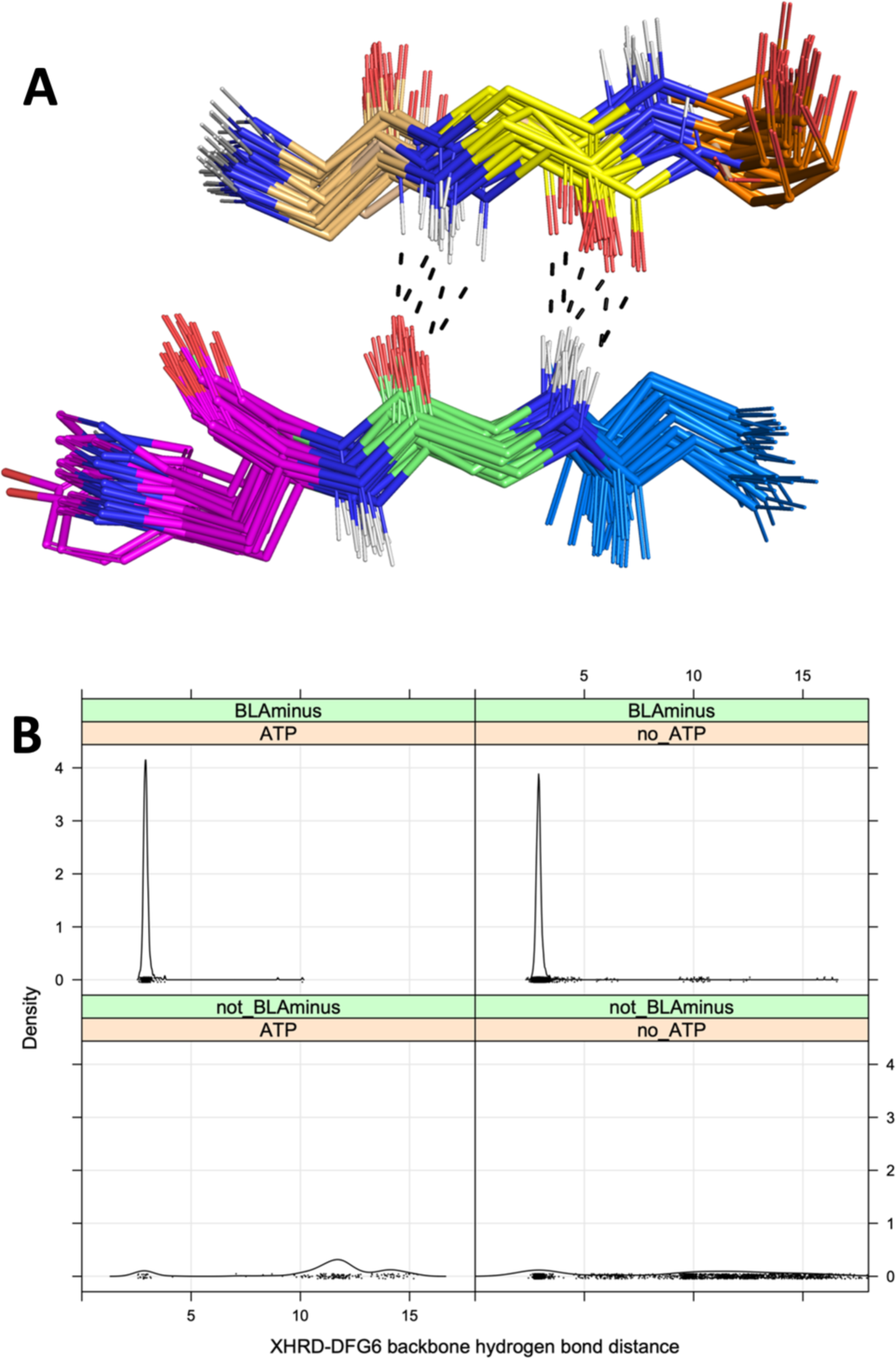
Interactions of residue 6 of the activation loop and the residue before the HRD motif (“XHRD”) **A.** Beta bridge hydrogen bonds between DFG6 and XHRD residues in kinase-substrate complex structures listed in Table 1. The carbon atoms are colored as follows: DFG5 (gold), DFG6 (yellow), DFG7 (orange), Xxhrd (blue), xXhrd (green), xxHrd (magenta, including the side chain, which is sometimes Tyr). Oxygen atoms are in red, hydrogen atoms are in white (modeled with PyMol), and nitrogen atoms are in blue. Hydrogen bonds in a few selected structures are marked with dashes. **B.** Distribution of the XHRD-DFG6 backbone-backbone hydrogen bond distance of ATP bound and unbound structures in the BLAminus and other states. The distance plotted is the minimum of the N-O or O-N distances between these two residues. The DFG6 residue is identified with the last X in the DFGxxX sequence, where x is any amino acid.

**Table 2.**
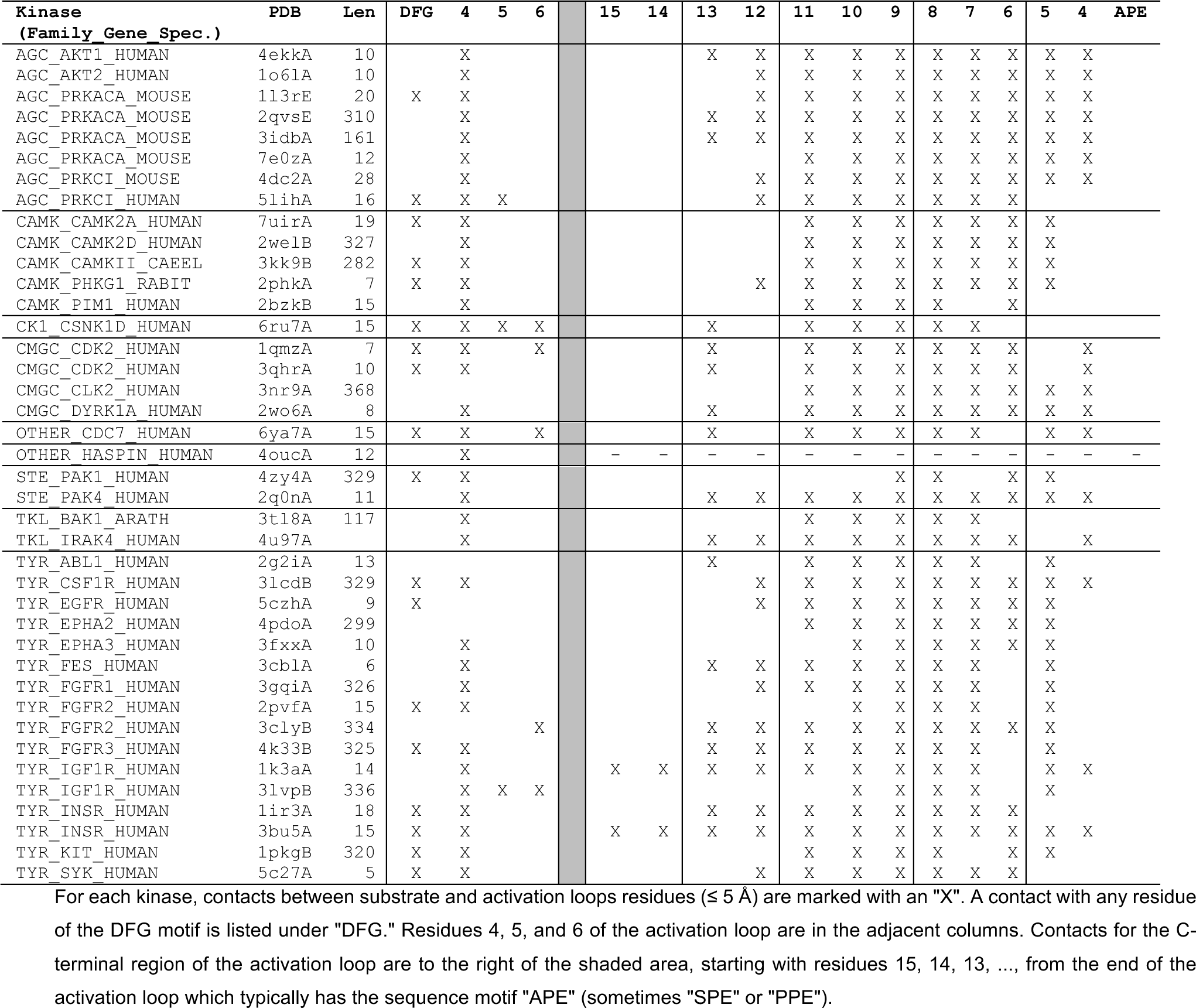
Contacts between activation loop residues and substrate.

Even without ATP, 97% of *BLAminus* structures contain the DFG6/XHRD hydrogen bond (Figure 6B, upper right). In the non-*BLAminus* state, only 24% of structures contain this hydrogen bond (Figure 6B, bottom panels).

### ActLoopCT

The conformation of the C-terminal end of the activation loop is critical for binding substrate. Most substrate-bound structures in Table 2 contain contacts between the substrate and residues 4-11 from the end of the activation loop, which ends in the sequence motif “APE.” From examination of the substrate-bound structures, we identified a contact that is consistent with substrate binding and which is absent in structures that likely block substrate binding: a contact (or near contact) between the APE9 Cα atom and the backbone carbonyl oxygen of the Arg residue in the HRD motif. This contact is shown in 23 non-TYR kinase structures from Table 2 in **Figure 7A**. The Cα-O distance is ≤ 4.2 Å in all of these structures.

**Figure 7.**
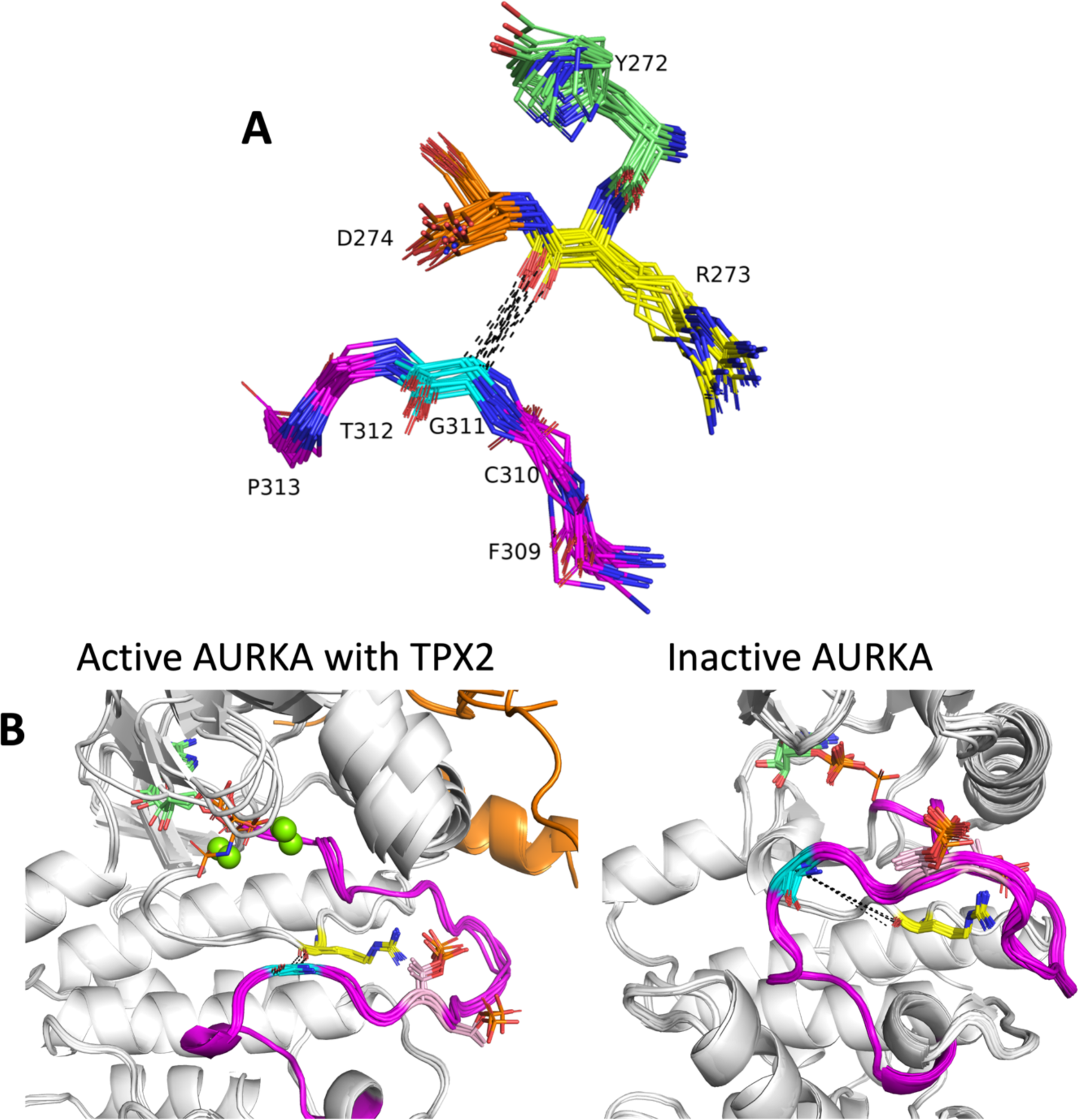
Role of the C-terminal segment of the activation loop in substrate binding. **A.** Contact between Ca of APE9 residue (XxxxxxAPE) and backbone carbonyl oxygen of Arg residue of HRD motif. His/Tyr (green), Arg (yellow), and Asp (orange) of HRD/YRD motif are shown in sticks including side chains, numbered according to AKT1 residues 272-274. Residues APE11 (AKT1 F309, magenta), APE10 (C310, magenta), APE9 (G311, cyan), APE8 (T312, magenta), APE7 (P313, magenta) are shown in sticks without side chains. Structures from the AGC, CAMK, CMGC, STE, and TKL kinases in Table 1 are shown. **B.** Two conformations of human AURKA. Left: Active structures with bound TPX2 (orange): PDB: 1ol5, 3e5a, 3ha6, 5lxm, 6vph. Right: Inactive structures without TPX2. PDB: 4dee, 5dt3, 5oro, 5oso, 6i2u, 6r49, 6r4d and others. Gly291 Cα (cyan sticks) is in contact with Arg255 backbone carbonyl O (yellow sticks) in the active structures (average distance 3.6 Å), while there is no contact in the inactive structures (average distance > 10 Å). The activation loop C-terminal region in the active structures resembles the structures of substrate-bound complexes in Figure 5, while the inactive structures would block substrate binding.

Aurora A kinase (AURKA) is a good example of the utility of these contacts. In the *BLAminus* state, there are two dominant conformations of the entire activation loop of AURKA. **Figure 7B** (left panel) shows five structures that contain these contacts. This comprises seven structures of AURKA with TPX2 (PDB: 1ol5, 3e5a, 3ha6, 5lxm, 6vpg). Two other structures bound with MYCN (PDB: 5g1x, 7ztl) are very similar (not shown). Both proteins are known to activate AURKA by binding to the N-terminal domain and the tip of the activation loop (Bayliss, Sardon et al. 2003, Richards, Burgess et al. 2016). Most *BLAminus* structures of AURKA, however, resemble the structures shown in Figure 7B (right panel). In these structures, the C-terminal end of the activation loop (APE6-APE10) deviates significantly from the TPX2- and MYCN-bound structures and from the structures of substrate bound kinases in the AGC and CAMK families. In the active structures, the Cα-O distances are about 3.6 Å, while in the inactive structures, the distance is more than 10 Å.

In Table 1, the APE9(Cα)-hRd(O) distance ranges from 3.4 to 4.2 Å in the substrate complexes in the Ser/Thr kinases (all families except TYR). This suggests that the Cα-O interaction is a CH-O hydrogen bond, which have been observed in proteins (Derewenda, Lee et al. 1995). In 271 of 355 non-TYR catalytic kinases, the APE9 residue is a glycine, which forms Cα-O hydrogen bonds more readily than other amino acids likely for steric reasons. In the substrate-bound TYR family kinases in Table 2, the APE9(Cα)-hRd(O) distance is longer and ranges from 6.5 to 7.4 Å.

We examined the distributions of this distance in ATP-bound and non-ATP structures in the *BLAminus* and other conformational states (**Figure 8**). For non-TYR kinases, the APE9-hRd distance is typically less than 6 Å in *BLAminus*/ATP-bound structures (Figure 8A, upper left panel), while the distance is much greater than 6 Å in a majority of non-*BLAminus* structures (Figure 8A, lower panels). As with the substrate bound structures, this distance is somewhat longer for *BLAminus*/ATP-bound structures of TYR kinases than for non-TYR kinases, ranging from 5 to 8 Å (Figure 8B, upper left panel), The large peak at 5 Å are all structures of FGFR2. For other TYR kinases, the distance is typically between 6 and 8 Å. One third of non-*BLAminus*, non-ATP-bound TYR kinase structures have an APE9-hRd distance greater than 8 Å (Figure 8B, lower right panel).

**Figure 8.**
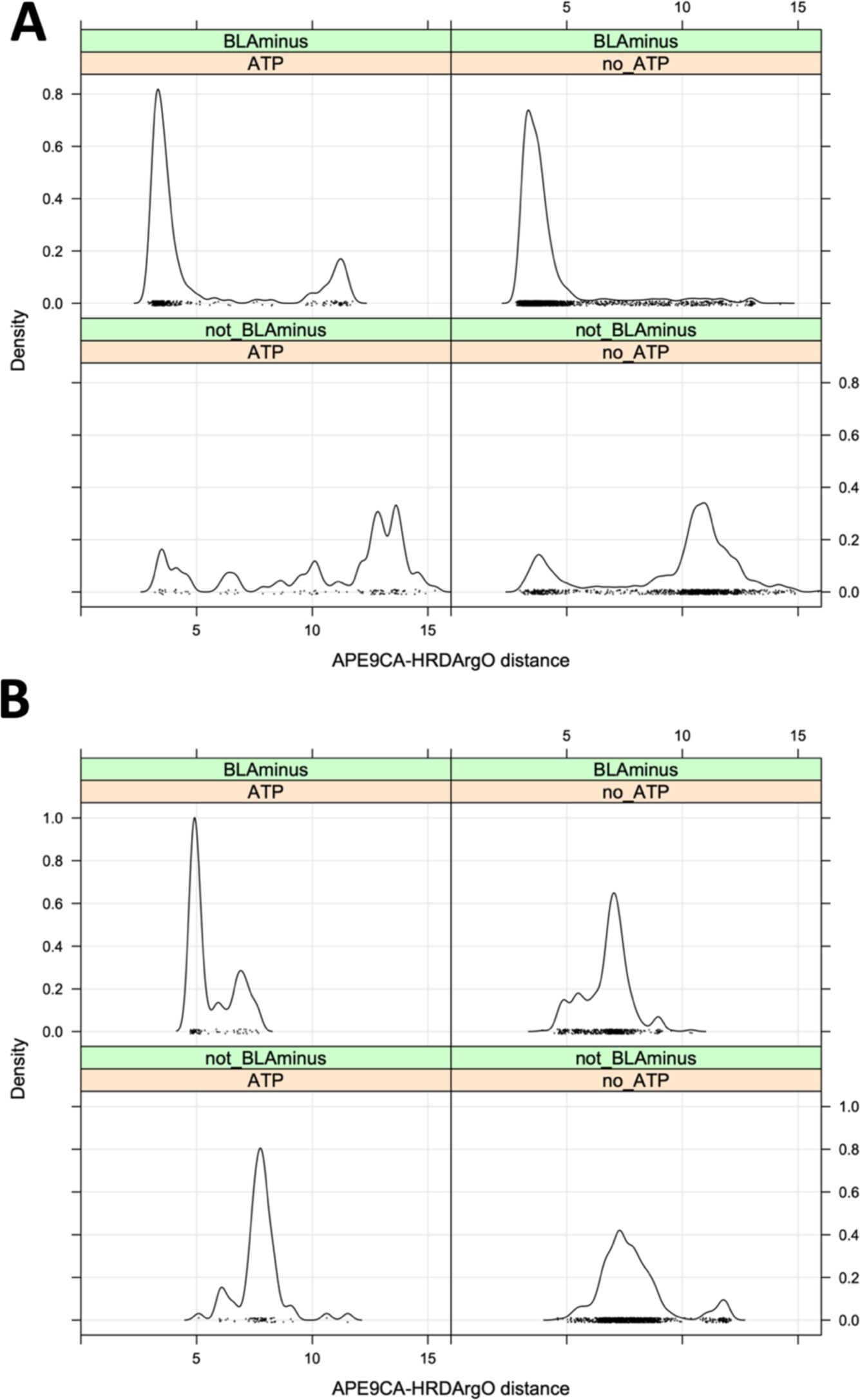
Criterion of residue 9 from the end of the activation loop (“APE9”) **A.** Distribution of the APE9-CA/hRd-O distance n ATP-bound and ATP-unbound structures. **B.** Distribution of the APE9-CA/hRd-O distance in ATP-bound and ATP-unbound tyrosine kinase structures. The peak at 5 Å in ATP-bound/BLAminus structures are all structures of FGFR2. The slightly longer distances from up to 8 Å are more characteristic of active tyrosine kinase structures. Longer distances (over 10 Å) are observed in non-BLAminus/non-ATP-bound structures (lower right panel).

### Regulatory spine

Finally, we evaluated the utility of the regulatory spine for identifying active structures. The regulatory spine consists of four amino acids:

1. the His residue of the HRD motif. This residue is: His in 393 catalytic kinases; Tyr in 38 AGC kinases, CK1_CSNK1G1,2,3; OTHER_SBK2; TKL_LRRK2; Leu in OTHER_PKDCC, Phe in TKL_LRRK1.
2. the Phe residue of the DFG motif. This residue Phe except: Leu in 38 catalytic kinases; Tyr in 11 catalytic kinases; Trp in CMGC_CSNK2A1,2,3; Met in CMGC_CDK8,19; Val in OTHER_PBK.
3. the Glu4 residue (corresponding to CAMK_AURKA Q185), which is four positions after the conserved Glu of the N-termina domain salt bridge. In catalytic kinases, this residue is: Leu (243 kinases); Met (114); Tyr (22); His (18); Phe (10); Ile (8); Gln (6); Cys (5); Val (4); Gly (3); Asn (2); Ser (2); Ala (2); Thr (1).
4. a usually hydrophobic residue just before the β4 strand, corresponding to L196 in CAMK_AURKA. We define this as “HPN7,” which means the seventh residue from the conserved HPN motif (HPNxxx***X***), which occurs in the loop between the C helix and the β4 strand. In catalytic kinases: Leu (256); Phe (58); Tyr (50); Met (25); Val (16); Ile (15); Cys (7); Ala (6); Thr (3); Gln (1); Ser (1).

These four residues define three distances: Spine1 (HRD-His/DFG-Phe), Spine2 (DFG-Phe/Glu4), and Spine3 (Glu4/HPN7). When the residues are small or polar, there may not be a contact between the side chains and such a contact may not be necessary for constructing an active kinase structure. In **Supplementary Figure 1**, the distribution of Spine1, Spine2, and Spine3 are shown for ATP-bound and unbound structures in the *BLAminus* and other states. From all three plots, it can be observed that nearly all ATP-bound, *BLAminus* structures contain an intact spine. The only exceptions are the Spine2 distances in two ATP-bound structures of PAK4 (PDB:7S46, 7S47) in which the C-helix is twisted by about 45° starting at the residue before the salt-bridge glutamic acid (E366). This distorts the position of the M370 side chain, which forms the Spine2 distance with DFG-Phe’s side chain. It is not known whether this distortion makes these PAK4 structures inactive, since the position of the Glu4 residue does not directly mediate contacts with ATP or the substrate.

Distributions of maximum spine distances (across Spine1, Spine2, Spine3) for kinase structures with and without ATP and in the *BLAminus* and other states is shown in **Supplementary Figure 2**. As we described in our previous paper, a majority of structures in several DFGin conformational states (*BLAminus*, *ABAminus*, *BLBplus*, etc.) contain an intact spine (defined as having all three spine distances less than 5.0 Å). 96% of *BLAminus* structures contain an intact spine. Most of those with a broken spine occur because of the position of the HPN7 residue, which does not interact with the substrate or ATP. When we apply the 5 active criteria described in the previous sections (DFGin, *BLAminus*, Saltbridge-in, ActLoopNT-in, ActLoopCT-in), there are 3013 human catalytic kinase chains in the PDB that are “Active.” If we define a broken spine as a structure with one or more spine residue pairs with a distance greater than 6 Å, there are only 14 structures in this set with an unformed regulatory spine, all of them PAK4 with a twisted end of the C-helix. There are only 11 active structures with a spine distance between 5 and 6 Å. For the sake of simplicity, we therefore do not use the regulatory spine as a criterion for active structures.

### Active structures of catalytic kinases in the Protein Data Bank

From the considerations above, we define probable “Active” structures of kinases at those capable of binding ATP, Mg ions, and substrate, with the following criteria:

1. DFGin spatial state
2. *BLAminus* dihedral angle state
3. SaltBr-in state (Nζ/Oε distance < 3.6 Å)
4. ActLoopNT-in (DFG6-Xhrd backbone hydrogen bond < 3.6 Å)
5. ActLoopCT-in (APE9-Cα/hRd-O distance < 6 Å in non-TYR kinases and < 8 Å in TYR kinases)

We made certain exceptions to the criteria for some kinases. The salt bridge criterion is skipped for OTHER_WNK1, WNK2, WNK3, and WNK4 kinases (WNK - “With No Lysine”) and for TKL_MAP3K12 and TKL_MAP3K13. In the experimental structures of TKL_MAP3K12 (e.g., 5CEP), the residue equivalent to the salt bridge Glu is Asp161 and is turned outwards with a break in the alpha C-helix, which is shorter than that of other kinases. The AlphaFold2 models with all of Uniprot90 as the MSA sequence database reproduce this unusual feature even in *BLAminus* structures. The presence of the Asp makes the salt bridge less likely to form so we omitted it as a criterion for these two kinases. Finally, OTHER_HASPIN, OTHER_TP53RK, and OTHER_PKDCC do not have APE motifs (Modi and Dunbrack 2019), and do not fold into the same structures as the C-terminal regions of other kinases. Thus, there is no ActLoopCT requirement for these kinases.

We calculated the relevant data for all human protein kinase domain structures in the PDB. The results are shown in Table 3 for catalytic kinases. Of 437 catalytic kinase domains in the human proteome, only 155 (35.5%) have active structures in the PDB. Of these, only 130 have complete sets of coordinates for the backbone of the activation loop, comprising less than 30% of catalytic kinases in the human proteome. We therefore chose to see if we could use AlphaFold2 to produce active structures of all 437 catalytic typical kinase domains in the human proteome.

**Table 3.**
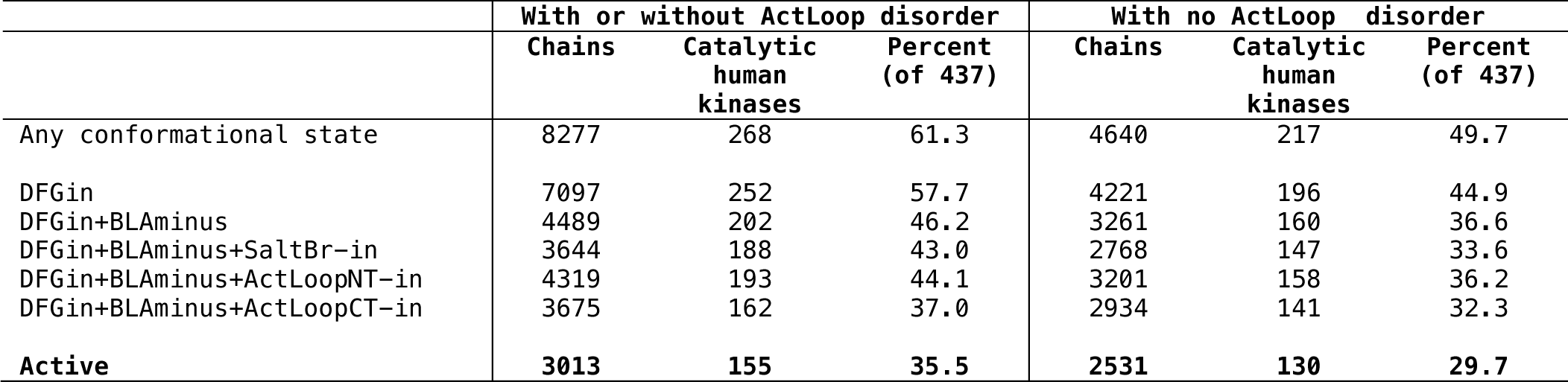
Classification of catalytic kinase domain structures in the PDB.

|  | With or without ActLoop disorder |  |  | With no ActLoop disorder |  |  |
| --- | --- | --- | --- | --- | --- | --- |
|  | Chains | Catalytic human kinases | Percent (of 437) | Chains | Catalytic human kinases | Percent (of 437) |
| Any conformational state | 8277 | 268 | 61.3 | 4640 | 217 | 49.7 |
| DFGin | 7097 | 252 | 57.7 | 4221 | 196 | 44.9 |
| DFGin+BLAminus | 4489 | 202 | 46.2 | 3261 | 160 | 36.6 |
| DFGin+BLAminus+SaltBr-in | 3644 | 188 | 43.0 | 2768 | 147 | 33.6 |
| DFGin+BLAminus+ActLoopNT-in | 4319 | 193 | 44.1 | 3201 | 158 | 36.2 |
| DFGin+BLAminus+ActLoopCT-in | 3675 | 162 | 37.0 | 2934 | 141 | 32.3 |
| **Active** | **3013** | **155** | **35.5** | **2531** | **130** | **29.7** |

### Generation of active models of catalytic protein kinase domains

To generate active models of the 437 human protein kinase domains, we created sequence sets for the multiple sequence alignments (MSAs) required by AlphaFold2 and template data sets in the active form. Sets of orthologous sequences (or near paralogs) for each kinase were created from UniProt such that each sequence in an ortholog set for a given kinase was greater than 50% identical to the target and aligns to at least 90% of the target kinase domain length with fewer than 10% gaps. Each ortholog set was filtered with CD-HIT so that no two sequences in a given set were more than 90% identical to each other. This was done to create diversity within the ortholog sets for each kinase. We also created “Family” sequence sets consisting of all the human kinase domains within each kinase family.

To create a template set, we identified all active structures of catalytic kinases in the PDB (including non-human kinases) using the criteria given above and selected two structures from different PDB entries (if available) with the largest number of coordinates for the activation loop residues (to select in favor of complete activation loops). If more than two structures were available with the same number of ordered residues in the activation loop, those with the highest resolution were selected. This resulted in a set we named “ActivePDB,” consisting of 165 kinase domains from 278 PDB entries.

We applied AlphaFold2 to all 437 human catalytic kinase domains, using the ortholog and family sequence sets and the ActivePDB template set. Different depths of the sequence alignment were utilized ranging from 5 sequences to 90 sequences. Only two of AlphaFold2’s five models (“model 1” and “model 2”) utilize templates, so only these models were run when templates were included in the calculations. If a structure of a target sequence was present in the template data set, it was removed when predicting the structure of the target. The models were relaxed with AMBER and the standard AlphaFold2 protocol, and we assessed the activity state of both the unrelaxed and relaxed models. In many cases, hydrogen bonds that were broken in the unrelaxed models were formed properly in the relaxed models.

We downloaded structures of all 437 kinases from the EBI website of AlphaFold2 models. The EBI site contains only one model per protein. Only 208 of the 437 kinase domains (48%) contain active structures within this set. When we ran all five models within AlphaFold2 with default parameters (with and without templates, Uniprot90 as the sequence database), we obtained active models of 281 catalytic kinases (out of 437) using the PDB70 template database and active models of 298 catalytic kinases using no templates. By comparison, under different conditions, using the ActivePDB templates and Distillation templates, ortholog and family sequence databases, and different MSA depths, we obtained between 371 and 421 active kinases (**Figure 9**) for each setup, depending on the input template and MSA data sources.

**Figure 9.**
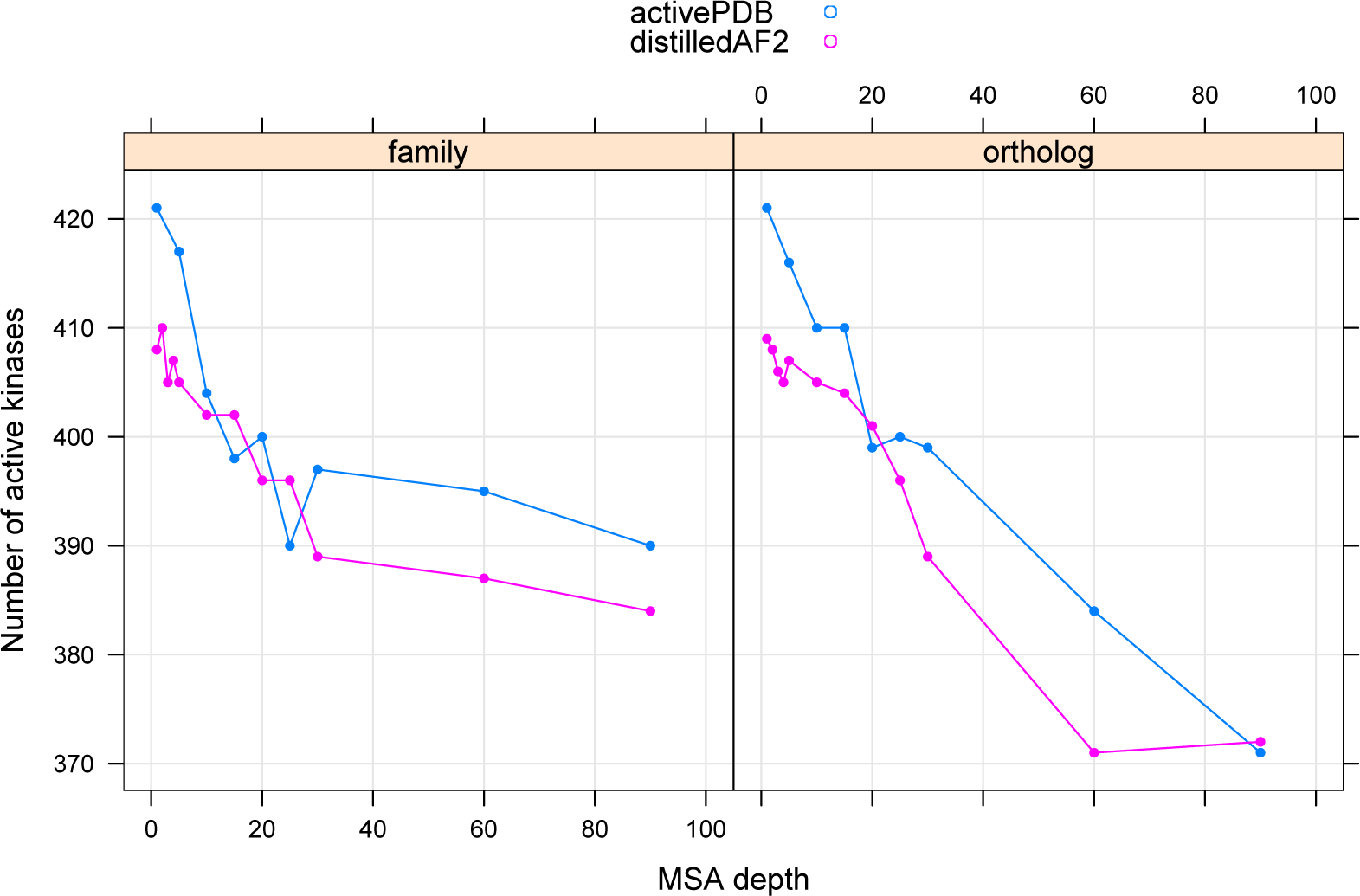
Number of catalytic kinases with active models produced by different template data sources (activePDB and distilledAF2) and different sequence sources (family kinases and orthologous kinases for each target).

No one set of inputs (MSA source, MSA depth, template database) produces active models of all 437 catalytic targets but combining the approximately 200 models from the different sets for each kinase achieved active models of 435 out of 437 targets. For two kinases, we needed special procedures. For the second kinase domain of obscurin (CAMK_OBSCN-2), the C-terminal segment of the activation loop made an α helix of residues 7825-7829 in all models that blocked access to the substrate binding site. We made the mutation D7929G (residue APE9, which is conserved as Gly in 73 out of 83 catalytic CAMK kinases) which helped to unfold this helix. It is possible that OBSCN-2 is a pseudokinase.

For LMTK2, all AlphaFold2 models formed a folded activation loop containing a strand-turn-strand motif that would be inconsistent with substrate binding. This structure forms in many DFGout structures of TYR kinase family members. We added additional distillation templates to the distilled AF2 template set of active structures of TYR_AATK (also known as LMTK1) and TYR_LMTK3. This produced active models of LMTK2 with very shallow sequence alignments (1-3 sequences from the ortholog data set). It also has an asparagine residue in place of the C-helix glutamic acid residue of the N-terminal domain salt bridge. However, LMTK2 has been shown to phosphorylate CFTR and other substrates involved in neuronal activity (Luz, Cihil et al. 2014).

In Table 4, we show the number of active kinase domains produced by different combinations of template database and MSA source summed over the MSA depths run for each combination shown in Figure 9. Using all the models with Uniref90 sequences produced active models of only 308 kinase domains. The ActivePDB template set plus the Family models and Ortholog models combined produced active models of 435 kinases. The only two kinases that required the distillation templates were LMTK2 and LMTK3; they only formed active models with 5 or fewer sequences from the ortholog set. As noted above, LMTK2 required the LMTK3 model, effectively a redistillation template. However, the quality of models in terms of pLDDTs of the activation loop is improved by including the distillation set models, as we show in the next section.

**Table 4.**
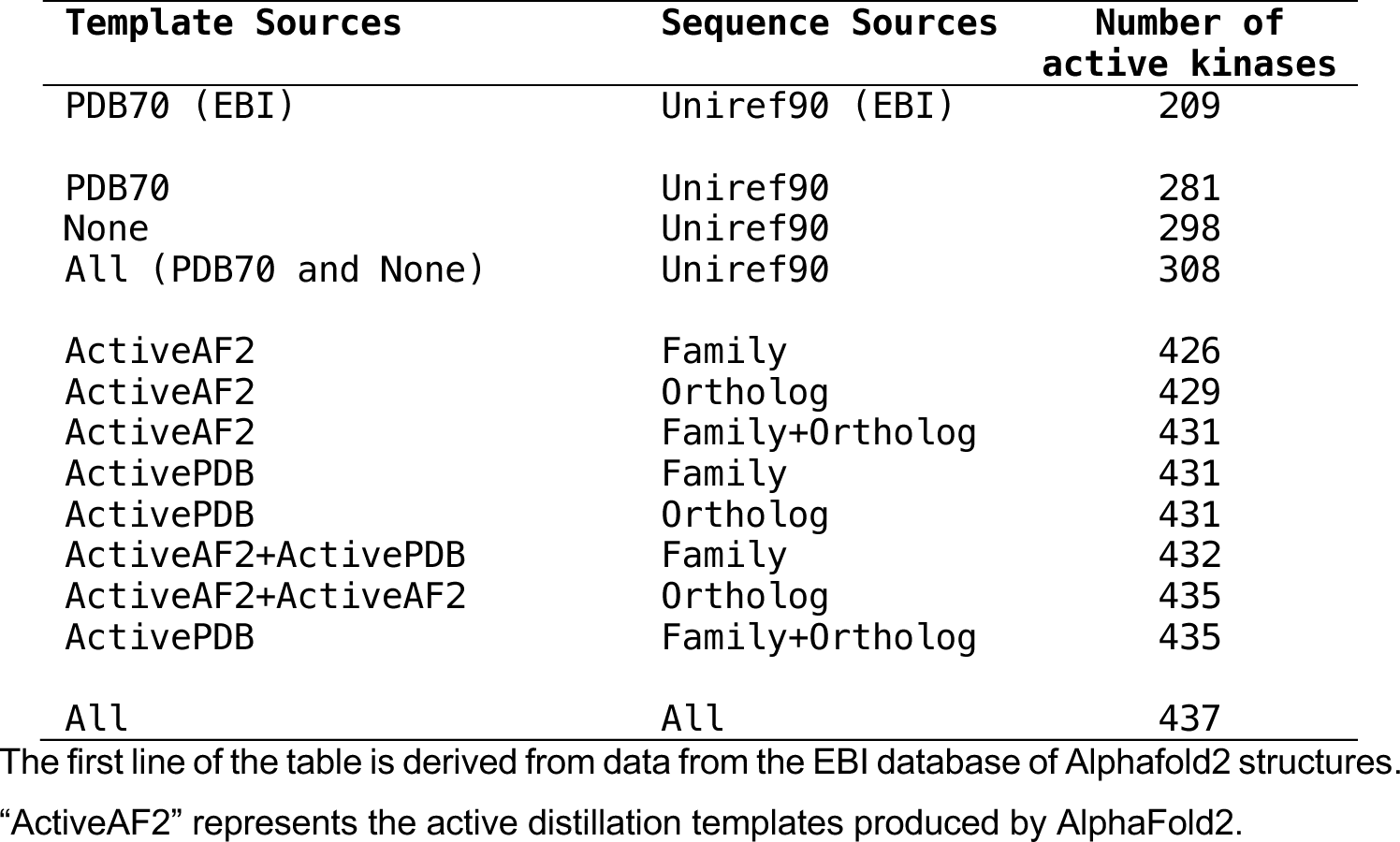
Number of active catalytic kinase domains (out of 437) produced by different Template and Sequence data sources.

### Picking the best model with pLDDTs scores of the activation loop

The structures of substrate-bound kinases (Figure 5) show that in active kinases, the activation loop is generally situated against the kinase domain, extending from the DFG motif towards the right-edge of the kinase domain (as generally oriented and shown in Figure 5). It then turns around and moves leftward and concludes in the APE motif, roughly below the DFG motif. This open U shape is characteristic of substrate-bound structures and of AlphaFold2 models produced by our pipeline. Dozens of examples of active AlphaFold2 models produced in this study are shown in **Figure 10** for the AGC, CMGC, STE, and TYR kinase families.

**Figure 10.**
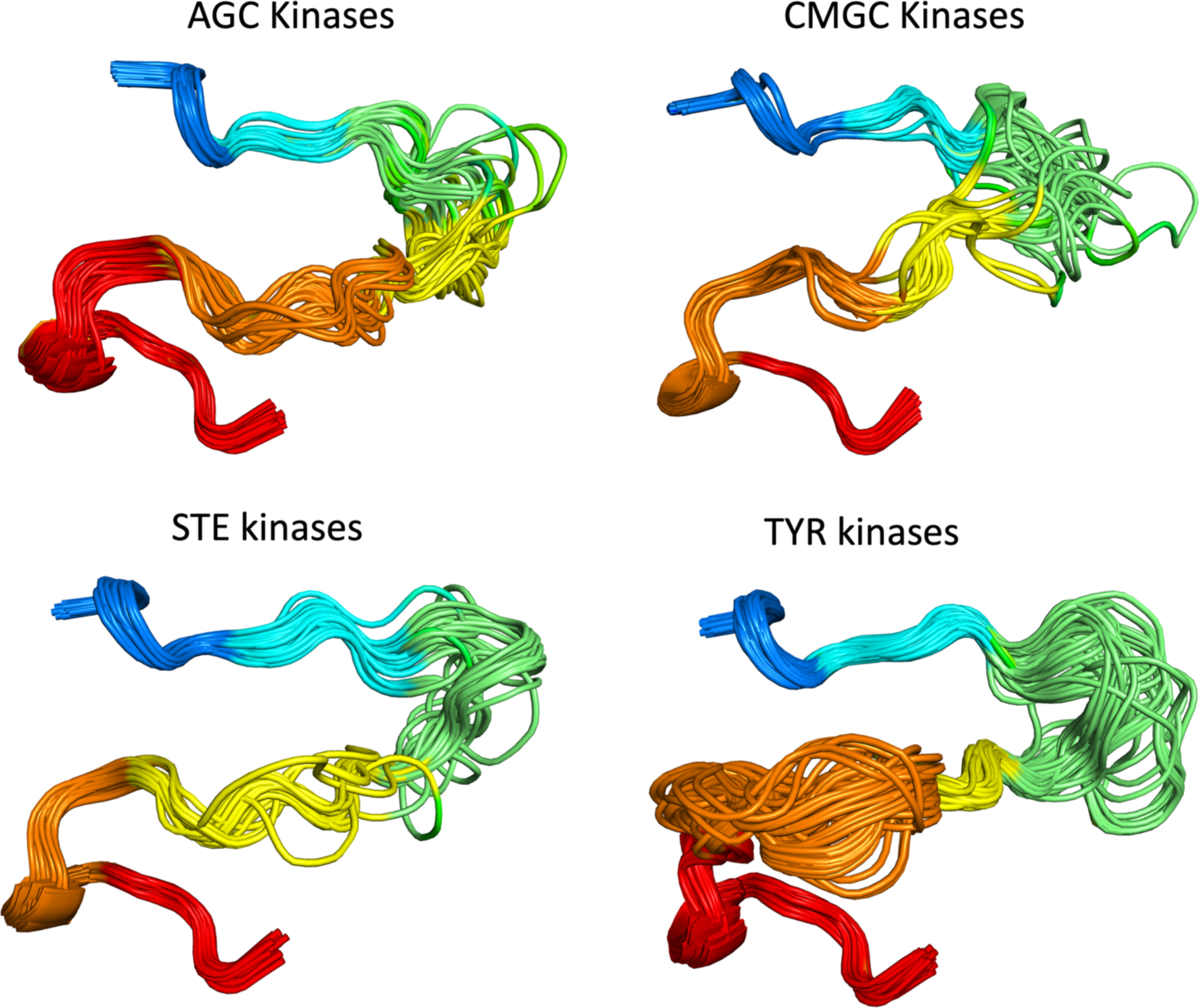
Examples of AlphaFold2 models of the active forms of 51 AGC kinases, 65 CMGC kinases, 31 STE kinases, and 77 TYR kinase. For clarity, some structures with long disordered regions within the activation loop are not shown in each family.

To benchmark the behavior of our pipeline in modeling active structures of catalytic kinases, we first compared the collection of AlphaFold2 models for the 22 kinases listed in Table 1 that have complete activation loops within their experimental structures. When the same kinase is listed more than once in Table 1, we picked a single example since the activation loop structures were all very similar (<0.5 Å RMSD).

These experimental structures contain substrates so are likely to be one (of possibly several) substrate-binding-capable conformations of the activation loop of each kinase. Both experimental and computed structures that pass our “Active” tests still exhibit some heterogeneity of the structure of the activation loop, especially for residues far from the beginning or the end of the loop. This may be natural structural variation and it is possible or even likely that multiple conformations are compatible with substrate phosphorylation. In any case, we explored the ability of the pLDDT values of the activation loop to pick out good models that pass our “Active” tests described above. We also wanted to know if the distillation templates produced better models of active structures in some cases.

In **Figure 11**, we show scatterplots of RMSD vs pLDDT of the activation loop for these 22 kinases. The results demonstrate that for most of the kinases, the highest pLDDT for the activation loop (defined as the minimum pLDDT value over the activation loop residues in the model) also produced the best or very close to the best RMSD to the structures listed in Table 1. The distillation templates (“ActiveAF2”) produced significantly better models than the ActivePDB templates for CMGC_CDK2 and CAMK_PIM1, and higher pLDDTs for most kinases. Thus, it seems likely that the extra sampling with the distillation templates may produce better models or more confident models for active structures of kinases.

**Figure 11.**
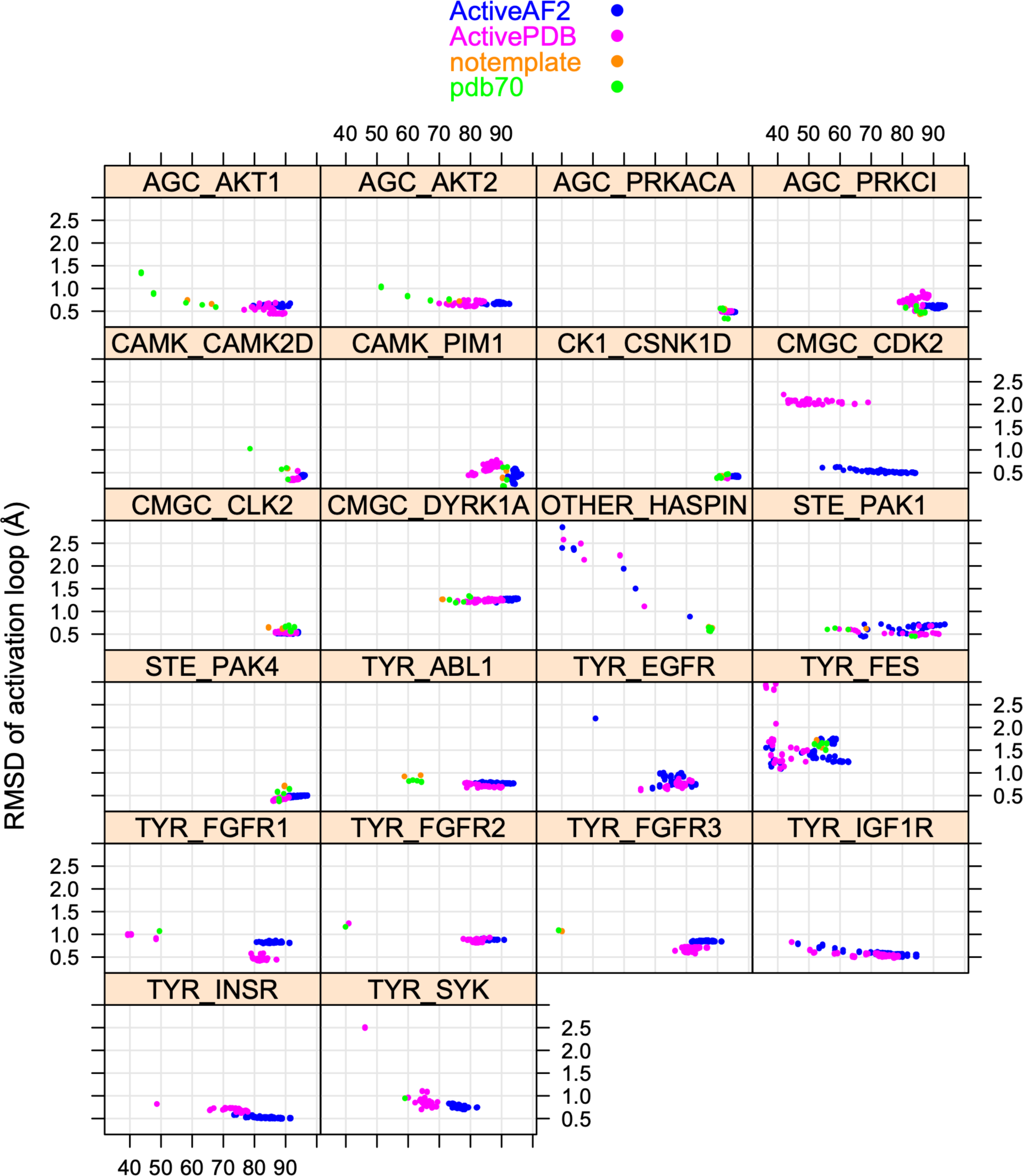
RMSD values for 22 substrate-bound structures from Table 1 versus the minimum value of pLDDT across the activation loop of each model. Models from different template data sets are shown in different colors: active structures from the PDB (“ActivePDB”), active models from AlphaFold2 (“ActiveAF2”), no templates provided to AF2 (“notemplate”), and AlphaFold2’s default template database (“PDB70”).

To extend the benchmark, we picked out at least one structure for each of the 130 human catalytic kinases with active structures in the PDB with complete activation loops. When all or almost all of the active structures for a particular kinase were similar (except for perhaps a few outliers), we picked out only one structure as a representative. When more than one conformation was represented in multiple PDB entries, we picked out a representative from each, labeling them “conf1,” “conf2,” etc. The structures labeled “conf1” were generally those that most closely resembled the substrate-bound structures in Table 1. The distribution of RMSD for the highest scoring models (highest min(pLDDT over the activation loop)) and the distribution of min_pLDDT values for the conf1 structures are shown in **Figure 12**. The results show that 104 (80%) of the 130 kinases are represented by a model with less than 1 Å backbone atom RMSD (N,Cα,C,O) over the whole activation loop (after superposition of the C-terminal domains of each kinase). A total of 117 (90%) are less than 2.0 Å.

**Figure 12.**
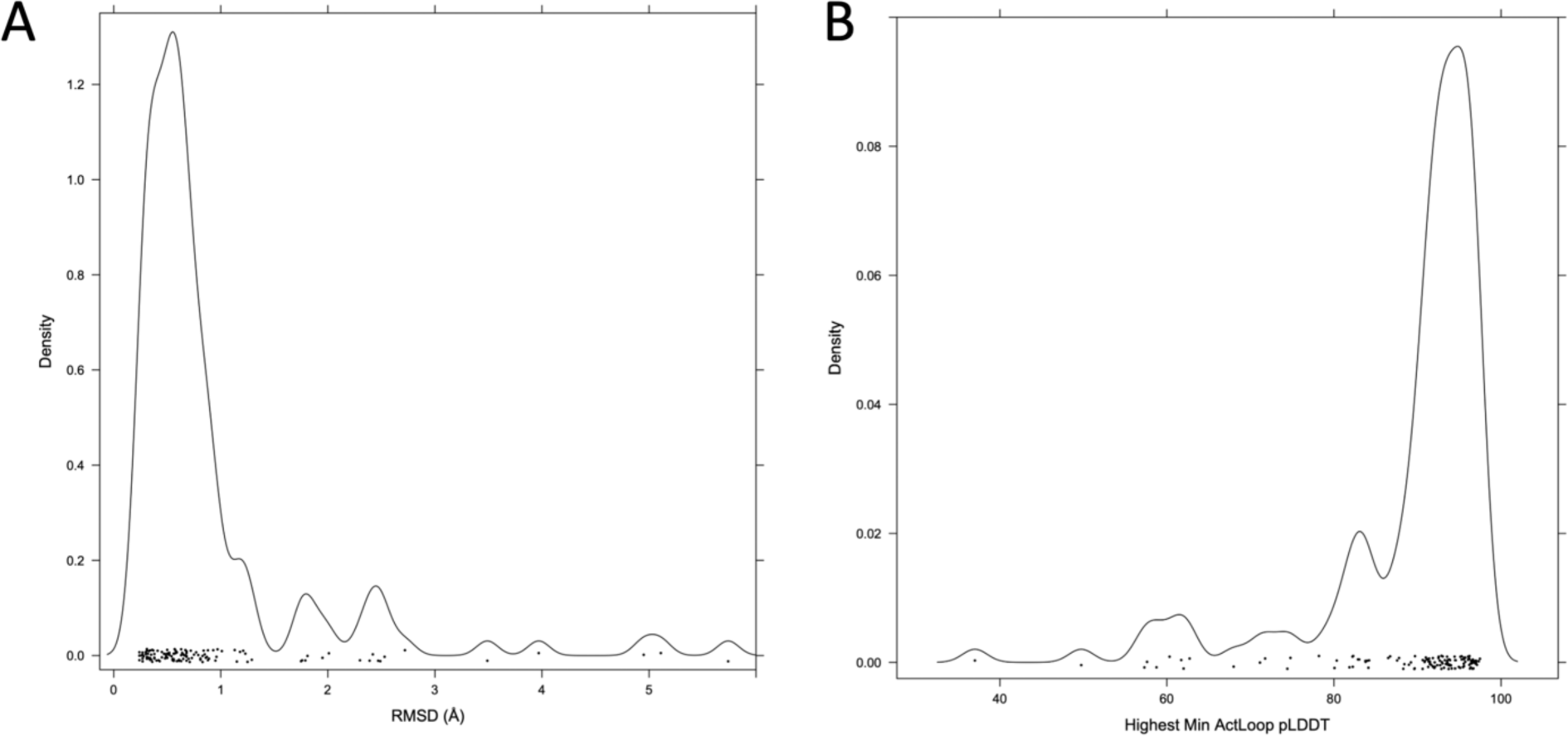
Benchmark of 130 active kinases in the PDB with full coordinates for the activation loop. **A.** Distribution of RMSD values (with kernel density estimate) for the top scoring model (minimum value of the pLDDT across the activation loop). **B.** Distribution of min_pLDDT values for these 130 models.

We can show that when multiple conformations of the activation loop of a given kinase are considered “Active”, our AlphaFold2 models are generally close to one of them, and this structure most closely resembles the substrate-bound structures in Table 1. For example, by visually clustering the structures of human CDK2 that pass our active-kinase criteria, we identified four predominant conformations (**Figure 13A**). The conf1 benchmark structure (PDB: 1QMZA) is a substrate-bound structure listed in Table 1. PDB structures very similar to the conf1 structure are also the only ones that are phosphorylated on residue T160 in the activation loop. For conformations conf2 (2BZKA), conf3 (5UQ1A), and conf4 (1FINA), the closest structures among the AlphaFold2 models have RMSD of 2.59 Å, 1.09 Å, and 2.78 Å respectively, all with min_pLDDT of less than 50.0. This contrasts with the best model of conf1, which has an RMSD of 0.48 Å to PDB:1QMZA and min_pLDDT of 84.2.

**Figure 13.**
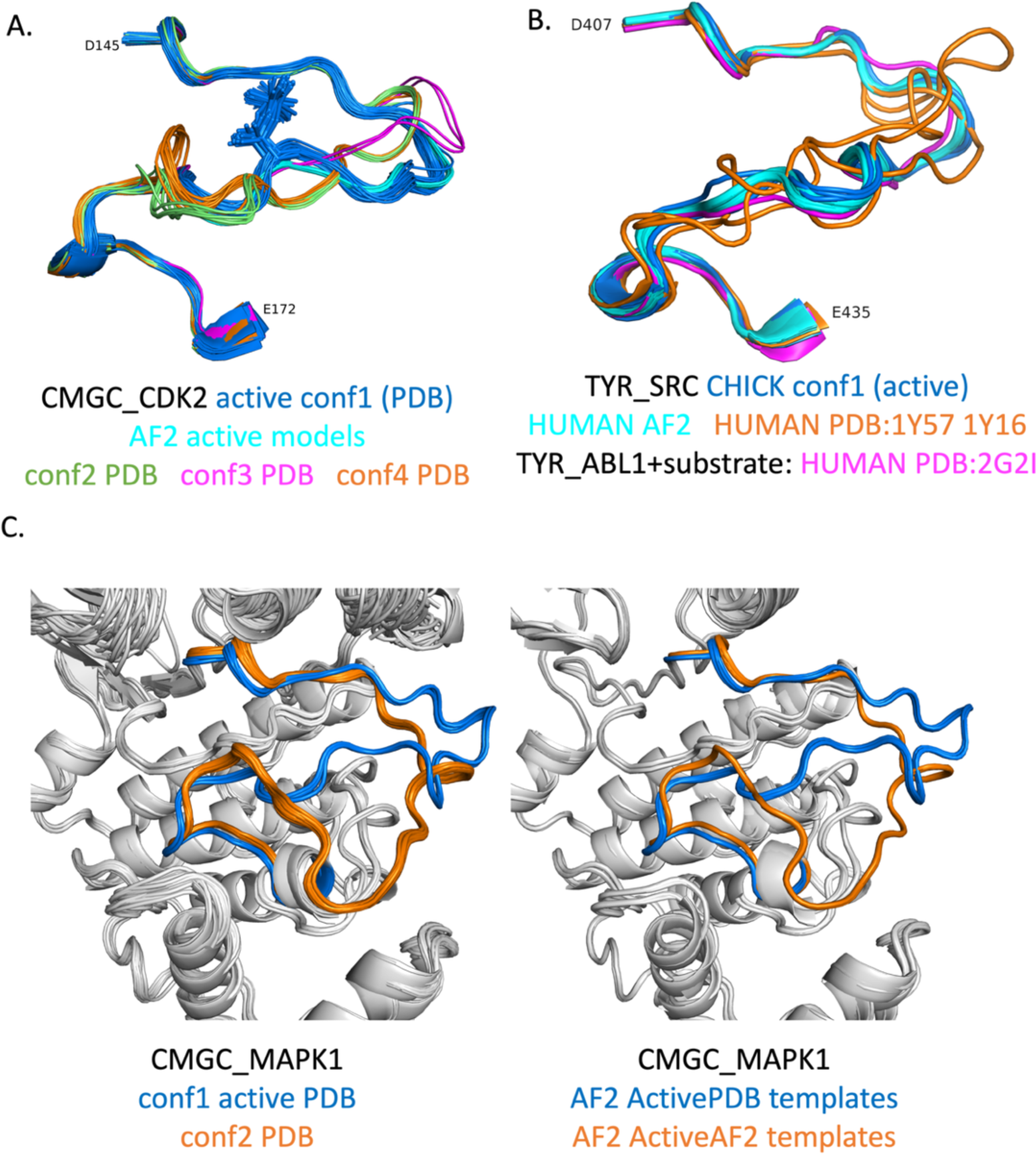
Structures of activation loops from the benchmark and corresponding AF2 models. **A**. CMGC_CDK2 has four dominant conformations among structures that pass our “Active” criteria in the PDB. The Conformation 1 cluster contains the substrate-bound structures listed in Table 1, and is also the only cluster that contains phosphorylated activation loops. The AlphaFold2 models most closely resemble Conformation 1. **B**.For TYR_SRC, we used a chicken SRC structure (PDB: 3DQW) as the benchmark structure since it most closely resembled substrate0bound structures of other TYR kinases such as ABL1 (e.g., PDB:2G2I in Table 1). The human structures which pass our criteria (PDB: 1Y57 and 1Y16 in orange) are quite different in much of the activation loop away from the first few and last few residues of the DFG…APE sequence. PDB: 1Y57 is often used as the basis of molecular dynamics simulations of “Active SRC” even though it is not likely the substrate- binding conformation. **C**. CMGC_MAPK1 has two dominant conformations in the PDB (left panel), one of which resembles substrate- bound structures in Table 1 (“conf1”) while the other has a bulge towards the C-terminus of the activation loop. AlphaFold2 reproduces both conformations (right panel), conf1 from ActivePDB templates and conf2 from ActiveAF2 (“distillation”) templates.

SRC presents another interesting example (**Figure 13B**). The human SRC structures which are “Active” by our criteria and contain fully ordered activation loops (PDB: 1Y57A (green in Fig 16B), 1Y16A, 1Y16B (orange in Fig 13B)) do not resemble substrate-bound structures of TYR kinases in Table 1, such as ABL1 (PDB:2G2I). However, there is a structure of chicken SRC in the PDB (PDB 3DQW, chains A-C, blue in Figure 13B) that is quite similar to the ABL1 (PDB 2G2I) structure listed in Table 1 (Figure 13B, magenta). The chicken SRC sequence differs by only two amino acids in the kinase domain from human SRC, neither of which is in the activation loop. Thus, there is no substrate-binding-capable structure of the SRC kinase domain annotated as human in the PDB, while the chicken SRC structure (PDB: 3DQW, chain A) presents a suitable model of active SRC. This is an important observation because an inactive structure of human SRC that is likely incapable of binding substrate (PDB: 1Y57) is often used as the basis of molecular dynamics simulations of the *active* protein (Foda, Shan et al. 2015, Fajer, Meng et al. 2017, Joshi, Burton et al. 2020). In the benchmark of 130 active structures, we replaced the structure of human SRC (PDB: 1Y57A) with that of chicken SRC (PDB: 3DQWA).

MAPK1 has two main conformations in our benchmark derived from the PDB (Figure 13C). One of them resembles the substrate-binding structures in Table 1 (Figure 13C, blue, left panel). The other has a bulge in the activation loop in the C-terminal half that places the activation loop over the APE motif and in contact with the G-helix (Figure 13C, orange, left panel). AlphaFold2 reproduces both of these structures almost exactly (Figure 13C, right panel) with RMSD of ∼0.3 Å in both cases. The active models are produced by the ActivePDB templates, while the alternate conformation models are produced by the ActiveAF2 distillation templates. The distillation templates included a structure of the closely related CMGC_MAPK3, which also has the same bulge as the orange structures in Figure 13C. The benchmark structure of MAPK3 (PDB: 4QTBA) in fact has the same bulge. However, the highest scoring AlphaFold2 model resembles substrate bound structures with an RMSD of 5.1 Å to the bulged benchmark structure 4QTBA, and is one of the RMSD outliers in Figure 12A. We believe this model more accurately reflects the likely substrate-binding conformation of MAPK3.

In addition to CMGC_MAPK1 and CMGC_MAPK3 just discussed, there are 11 other kinases where the highest pLDDT models have more than 2 Å RMSD to the structural representative we chose These kinases are: CMGC_HIPK3 (2.01 Å RMSD), CMGC_MAPK7 (5.74 Å), OTHER_BUB1 (3.49 Å), OTHER_WNK3 (2.72 Å), STE_MAP3K14 (3.97 Å), STE_MAP3K5 (2.42 Å), STE_STK3 (2.30 Å), STE_TNIK (2.47 Å), TKL_ACVR1 (2.49 Å), TKL_ACVR2B (2.39 Å), and TKL_BRAF (2.53 Å). These are analyzed and discussed in **Supplementary Figure 3** and **Supplementary Figure 4**. In some cases, the model and PDB structure only differed in the outermost residues of the activation loop (from the DFG and APE motifs). This occurred for STE_STK3, STE_TNIK, TKL_ACVR2B, and OTHER_WNK3 for example. In TKL_ACVR2B, there is a change in position of residues 10-17 or a 30-residue activation loop, while residues 1-9 and 18-30 are very similar in the benchmark structure (2QLUA) and the AF2 models.

In some other cases, the AF2 structure appears capable of binding substrate while the PDB structure does not. This can be demonstrated by comparing the benchmark structure to that of closely related kinases in the PDB and in our AF2 models. For example, the CMGC_HIPK3 and CMGC_HIPK2 benchmark structures are quite different in the C-terminal region of the activation loop. The AF2 models of HIPK2 and HIPK3 closely resemble the HIPK2 experimental structure (PDB:7NCFA) but not the HIPK3 experimental structure (PDB: 7O7IA). The activation loop sequences of HIPK2 and HIPK3 are 96% identical (25 of 26 positions). Similarly, for TKL_ACVR1A, the PDB structure (6UNSA) blocks the active site, while the AF2 models resemble the TKL kinase BAK1 (PDB:3TL8A) from Table 1, which contains a substrate peptide.

For some other kinases, the AF2 models have poor pLDDT scores in the activation loop. This occurs for some kinases that are remotely related to other kinases in the human proteome or that have particularly long activation loops. For all three kinases in the RAF family (ARAF, BRAF, RAF1 or CRAF), the min_pLDDT scores for the activation loop are below 40. For BRAF, the top scoring AF2 models are not very similar to the benchmark structure with an RMSD of 2.53 Å (PDB:4MNEB, the only structure of BRAF with a complete activation loop that passes our “Active” criteria). It is unclear if this PDB structure is fully capable of binding substrates or whether the AF2 models are in fact better models of substrate- capable structures.

## DISCUSSION

We have developed a structural bioinformatics approach to identifying structures of typical protein kinases that are likely capable of binding ATP, metal ions, and substrates and catalyzing protein phosphorylation, which is involved in nearly all cellular processes in eukaryotes. We applied these criteria to experimental structures, which enabled us to develop a set of templates that could be used to model all 437 catalytic protein kinases in their active form with AlphaFold2. The same criteria enabled us to distinguish active structures among the models produced by AlphaFold2, which we cycled back into the protocol as templates for producing additional models with improved pLDDT scores. We refer to these as distillation templates, in analogy to the distillation models that the team at DeepMind used as additional training data for the original implementation of AlphaFold2. We demonstrated that the models with the highest values of pLDDT for the activation loop residues also most closely resemble substrate-bound structures of kinases in the PDB. In all, we generated approximately approximately 90,000 models to identify active model structures for all 437 catalytic human kinases.

While much attention has been given to the structure of the active site residues surrounding ATP, including the DFG motif and the N-terminal domain salt bridge, we examined 40 substrate-bound structures of protein kinases in the PDB to define criteria that ensure the presence of a substrate binding site necessary for the phosphorylation reaction. This is a far larger set of substrate-bound structures than has been previously analyzed, since it includes known autophosphorylation complexes contained within crystals (Xu, Malecka et al. 2015). In substrate-bound structures, the activation loop is extended away from the ATP binding site, lying against the surface of the kinase domain. To accomplish this, the activation loop interacts with the relatively fixed positions of residues in the catalytic loop in and around the HRD motif. This occurs near the N-terminus of the activation loop in a short beta sheet (beta6 + beta 9, (Kornev and Taylor 2010)) and can be identified by backbone-backbone hydrogen bonds of residue 6 of the activation loop with the residue that immediately precedes the HRD motif. It also occurs near the C-terminus of the activation loop where the Cα atom of residue 9 from the end of the loop makes a short contact with the backbone carbonyl of the Arg residue of the HRD motif. While other distances could also be used as criteria, we found that all substrate-bound structures in the PDB satisfy these two rules and that the vast majority of experimental and computed structures that satisfy these criteria appear to form a functional substrate binding site. For some kinases, there remains some conformational diversity of the activation loop among PDB structures after satisfying these criteria. It is likely that multiple conformations of the outer portion of the activation loop may be capable of phosphorylating substrates in some kinases. In other cases, some conformations that satisfy our criteria may block substrate binding.

Unfortunately, there does not seem to be a readily identifiable criterion that would be applicable across kinases to identify such situations. This phenomenon does seem to be rare. For example, MAPK1, MAPK3, and MAPK7 share an alternate conformation in experimental structures that would block substrate binding. AlphaFold2 produces these structures but also substrate-capable structures that resemble substrate-bound structures in the same CMGC family. These latter models are the ones we have made available in a set of models of active structures of all 437 catalytic typical protein kinases in the human proteome (http://dunbrack.fccc.edu/kincore/activemodels).

We believe our models will be useful in understanding the structural basis of kinase substrate specificity, since they place substrate binding residues in the activation loop and the active site of the kinase domain in suitable positions for catalysis. There remain challenges, however. We have found that AlphaFold-Multimer is in some cases capable of making models of substrate-bound structures of typical protein kinases when given a peptide substrate and Uniref90 as a sequence database. But it is not always able to do make an active model of the kinase activation loop without appropriate templates and shallow sequence alignments. But doing this sometimes disrupts its ability to place the substrate in the active site, probably due to the lack of sequence information for the substrate MSA. This will take additional study and implementation to develop a robust protocol that reliably makes models of kinase-substrate complexes from suitable choices of templates and multiple sequence alignments for AlphaFold-Multimer. This work is ongoing.

## METHODS

### Ortholog sequence sets

We first searched UniProt for Pfams PF00069 and PF07714 to collate a set of 1.68 million sequences in UniRef100 with typical protein kinase domains. For each of 437 catalytic kinase domain sequences from our earlier alignment of all human kinase domains (Modi and Dunbrack 2019), we used PSI-BLAST to get a list of the top 25,000 closest kinases to each human kinase domain. The queries used were 8 residues longer on each end of the kinase domain than our published alignment. The hit regions in the PSI-BLAST output were then filtered for sequences more than 50% identical to the query, coverage greater than 90% of the query length, and gap percentage in the alignment of less than 10%. We then applied CD-HIT (Fu, Niu et al. 2012) to create lists of orthologs (or close paralogs) with no more than 90% sequence identity to each other. These sequences were used as query databases in AlphaFold2 calculations.

### AlphaFold2

We used DeepMind’s advanced machine learning model, AlphaFold2, to predict the structures of human catalytic kinases. The code for AlphaFold2 was downloaded from DeepMind’s official GitHub repository (https://github.com/deepmind/alphafold). The computations were performed on workstations with NVIDIA GeForce GPUs (8, 12, or 24 Gbytes each). Each system was equipped with Linux (Ubuntu 20.04), CUDA11, Python 3.8, and TensorFlow 2.3.1.

### Data Input and Preparation

Three sets of sequence databases were used to create multiple sequence alignments: the default UniRef90 database, an additional kinase family-focused sequence database (all 496 human kinases in the human proteome, separated into each family), and a kinase orthologs-focused sequence database (described above). Templates for the calculations were obtained from the default PDB70 set, a curated selection of active PDB models identified through our criteria by Kincore (“ActivePDB”), and a distilled set of AlphaFold2 models that passed Kincore criteria with activation loop pLDDT scores of 60 or higher (“ActiveAF2” or “distilled”).

### Model Configuration and Implementation

Calculations with AlphaFold2 were conducted using the recommended configurations provided by DeepMind. The multiple sequence alignment was prepared using the hh-suite package (Steinegger, Meier et al. 2019) and subsequently fed into the model for structure prediction. When using templates, we used only AlphaFold2 models 1 and 2 since they utilize templates and the MSA data, while models 3, 4, and 5 do not use templates (Jumper, Evans et al. 2021) which was done by commenting out models 3-5 (lines 39-61 in /alphafold/model/config.py):

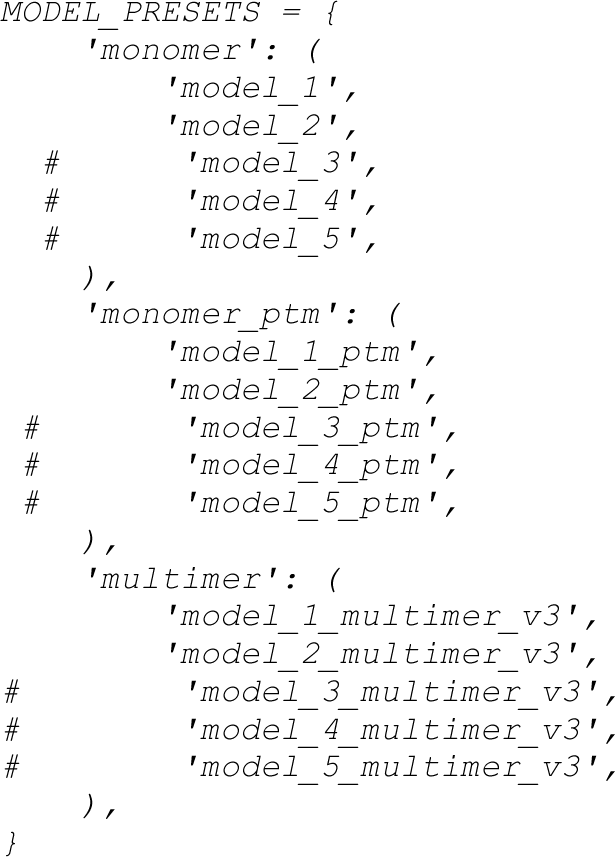

We ran AlphaFold2 with specific sequence data sets by replacing ./uniref90/uniref90.py with our sequence sets: Uniref90, Ortholog, and Family for the MSA building step and specific template data sets to predict protein structures. Template implementation consisted of two parts: .cif files of the structures in `./pdb_mmcif/mmcif_files` and their sequence data in the `./pdb70` folder. For each set of templates (PDB70, ActivePDB, ActiveAF2), the files need to be changed for AF2 to use the desired template set. When predicting the structure of a given target, any structures of the target in the PDB template sets were removed.

We introduced a variable `MSAlimit` that controls the number of sequences in the multiple sequence alignment used by AF2 model building by modifying the class `DataPipeline` (in /alphafold/data/pipeline.py). When AF2 has too many sequences in the MSA, it tends to ignore any templates provided to it. We also disabled other sequence databases like mgnify, bfd, small bfd, uniref30:

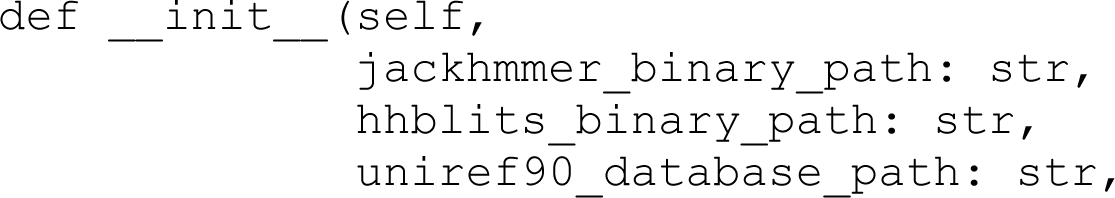
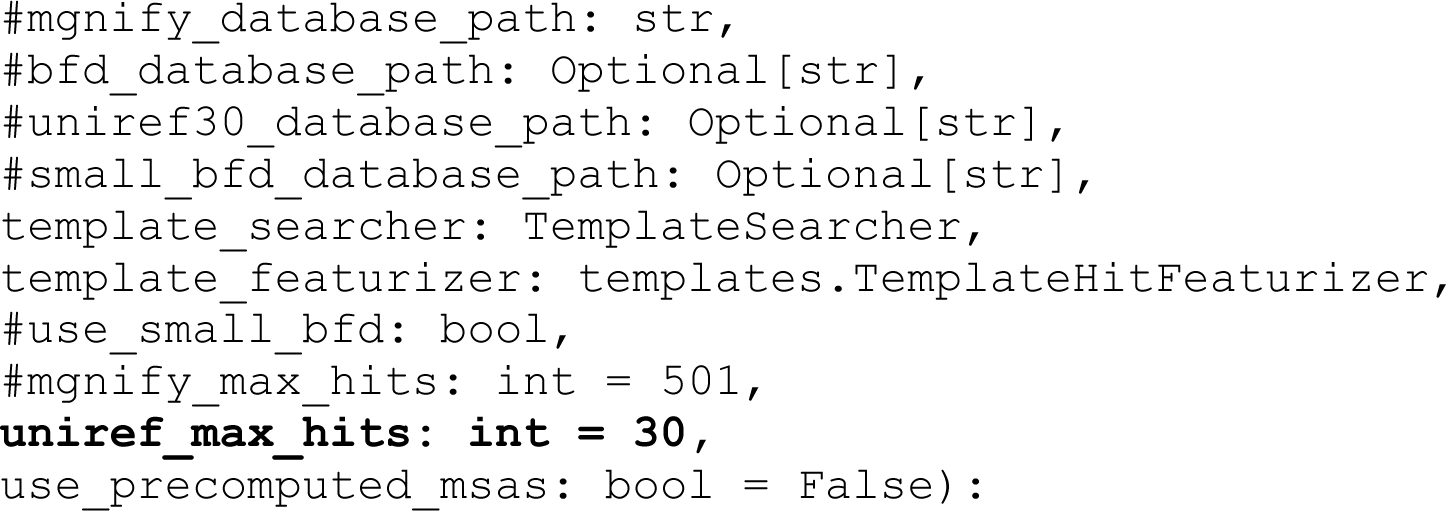

### Benchmarking

The predicted structures were validated by comparing them to the benchmark PDB structures of kinases. The validation process relied on the pLDDT score (Mariani, Biasini et al. 2013) and Root Mean Square Deviation (RMSD), measuring the average distance between atoms in the predicted and known structures. Model structures were aligned to benchmark structures with the program CE (Shindyalov and Bourne 1998) as implemented in PyMOL. The alignment was performed on the C-terminal domain of each structure. RMSD was measured for the activation loop backbone atoms (N, CA, C, O) after superposition of the C-terminal domains.

Two benchmarks were constructed. One contained substrate-bound structures from Table 1 with complete coordinates for the activation loop in the PDB structure (22 kinases). The other consisted of 170 structures of 130 with complete activation loops that passed our active criteria from the PDB. For some kinases, there were multiple conformations that passed our criteria. We labeled the structure that most closely resembled substrate-bound structures as “conf1” with the others labeled “conf2,”, “conf3,” etc.

### Update to Kincore Database and Website and Kincore_Standalone2

We have updated the Kincore database and website (http://dunbrack.fccc.edu/kincore) to include the additional active criteria defined in this study (Saltbridge, ActLoopNT, ActLoopCT) for all structures in the PDB. Each structure is marked “Active” or “Inactive” based on these criteria. We have also added the highest scoring AlphaFold2 active model for each of the human catalytic kinases to Kincore. These are labeled with the prefix “AF-” and the suffix “-K1” attached to the Uniprot Accession ID (e.g., the CAMK_AURKA model is AF-O14965-K1). The updated standalone program for assessing the conformational state of protein kinases, Kincore-standalone2, is available at https://github.com/DunbrackLab/Kincore-standalone2/.

### Data Availability and Reproducibility

To ensure the reproducibility of our study, all data, including input sequences, ortholog sequence sets, and predicted structures, are accessible at http://dunbrack.fccc.edu/kincore/activemodels.

## ACKNOWLEDGMENTS

This work was funded by NIH Grant R35 GM122517 (to RLD) and P30 CA006927 (to Fox Chase Cancer Center).

---

**Supplementary Figure 1.**
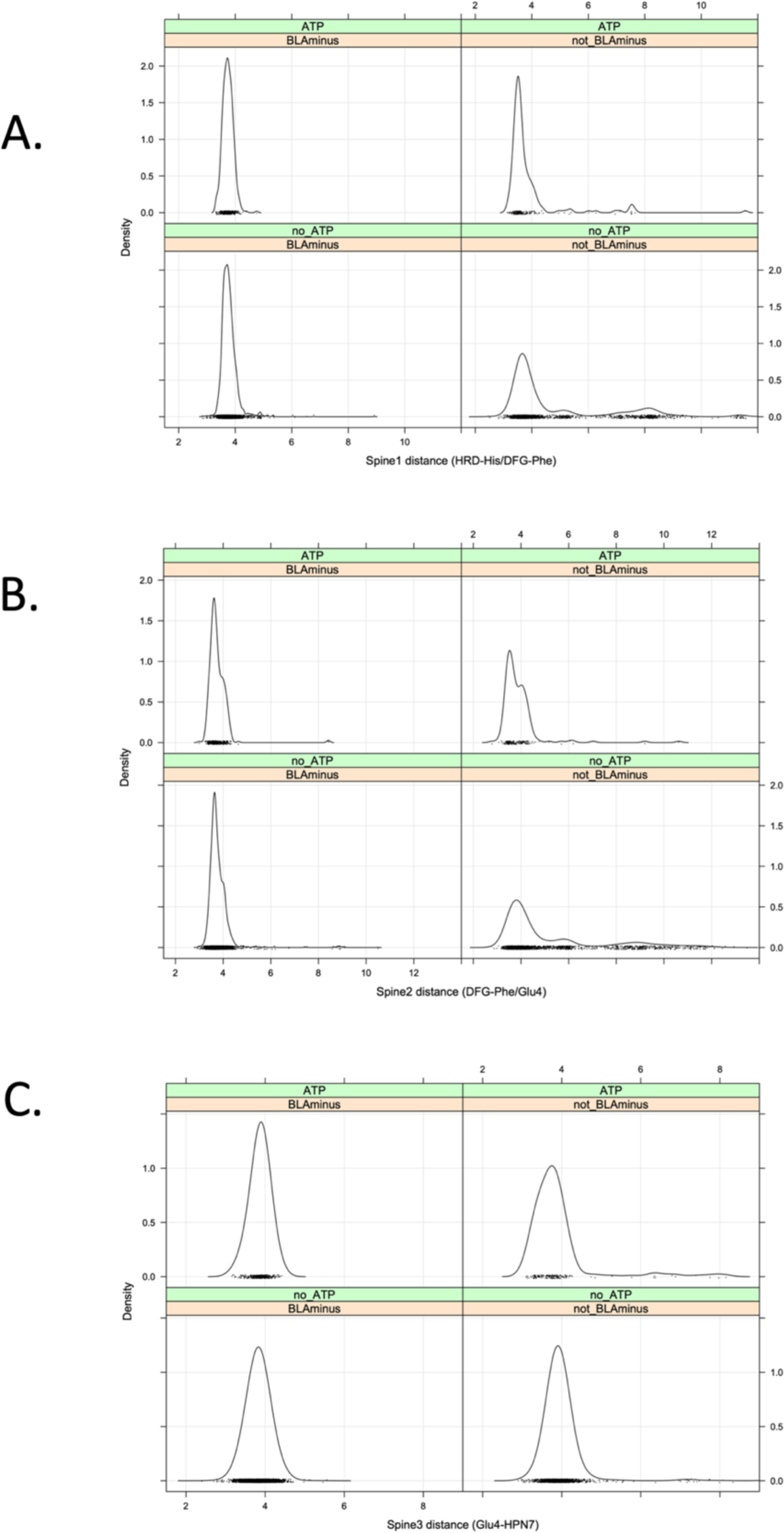
Distribution of the Spine1, Spine2, and Spine 3 distances. The Spine distances are defined as the closest distance among all side-chain atom pairs between the two residues.

**Supplementary Figure 2.**
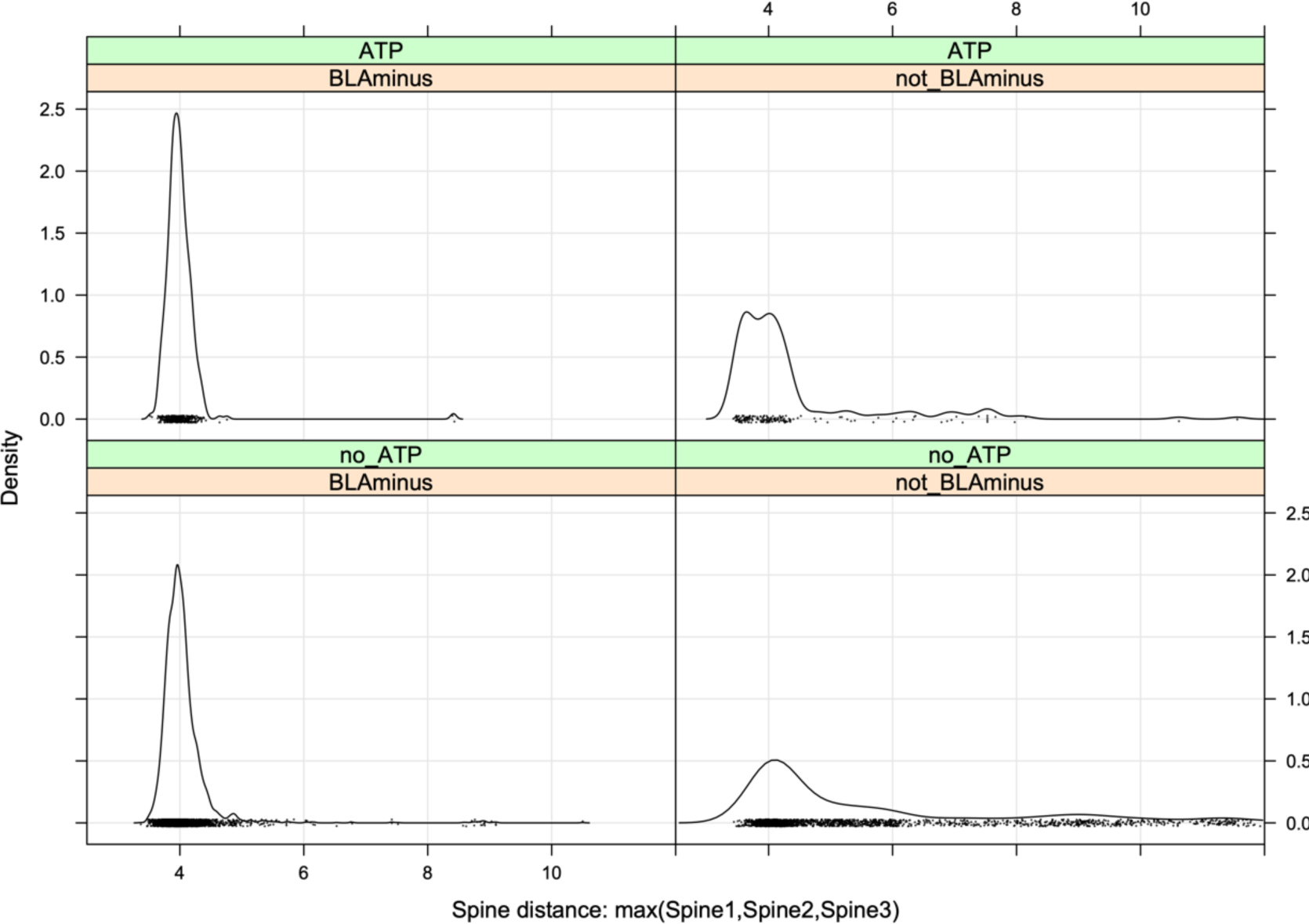
Distribution of the Spine distance, which is the maximum of Spine1, Spine2, Spine3 (Supplementary Figure 1) for each kinase structure.

**Supplementary Figure 3.**
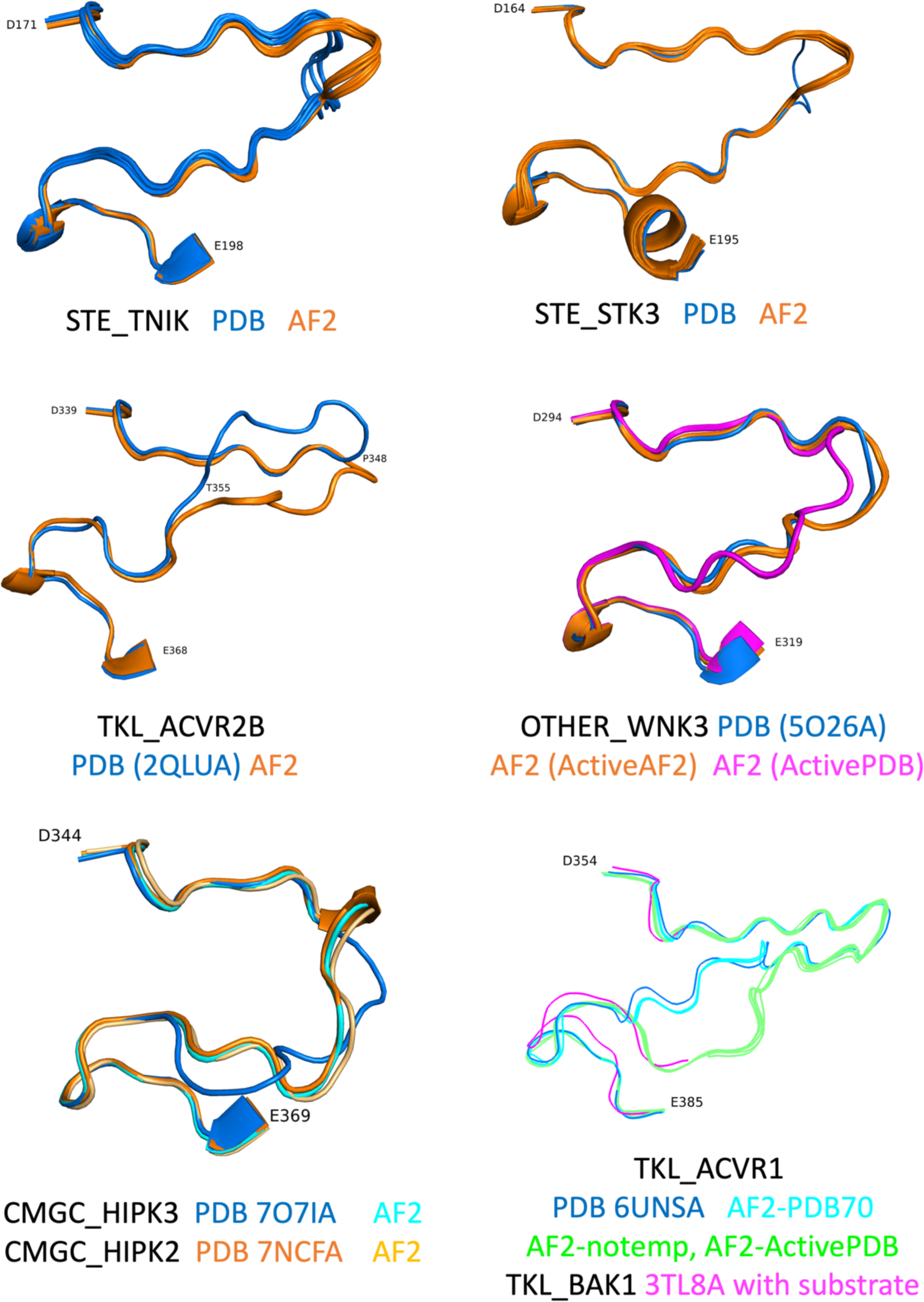
Benchmark structures with large RMSD to the best scoring AlphaFold2 models. STE_TNIK, STE_STK2, and TKL_ACVR2B show deviations between the PDB structures (blue) and AlphaFold2 models (orange) only in the outer regions of the activation loop. For OTHER_WNK3, the highest scoring pLDDT model is from an ActivePDB template (magenta) while an ActiveAF template model is much closer (orange) to the benchmark structure (PDB:5O26A, blue). For CMGC_HIPK3, the AF2 model more closely resembles an active PDB structure of HIPK2 (PDB: 7NCFA, orange) as well as an AF2 model of HIPK2 (light orange) than a PDB structure of HIPK3 itself (7O7IA, blue). The activation loop sequences are identical in 25 of 26 positions, and it is likely that AF2 is correct in modeling very similar conformations of the activation loop. The PDB structure of TKL_ACVR1 (PDB: 6UNSA. blue) resembles the AF2 model made from PDB70 (cyan), while the ActivePDB AF2 model and the no-template AF2 model more closely resembles a substrate-bound PDB structure of TKL_BAK1 from *Arabidopsis* (PDB: 3TL8, magenta, substrate not shown). It is likely that the ActivePDB AF2 model (green) is a substrate-binding structure, while the PDB structure is not.

**Supplementary Figure 4.**
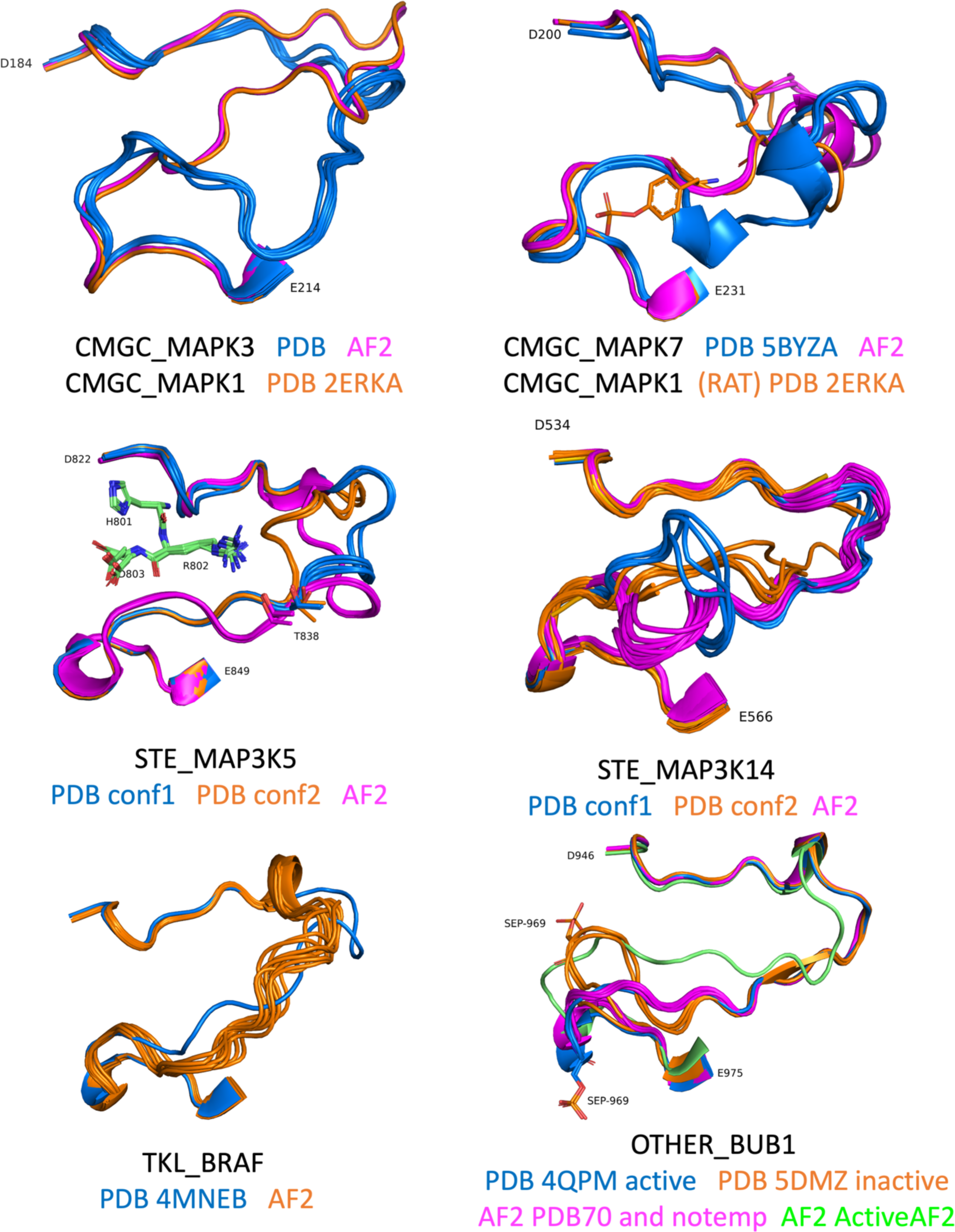
Benchmark structures with large RMSD to the best scoring AlphaFold2 models. CMGC_MAPK3 and CMGC_MAPK7 show the same bulge toward the C-terminal end of the activation loop that is present in CMGC_MAPK1 (Figure 13C, main paper) in PDB structures (blue). The AF2 structures (magenta) more closely resemble RAT CMGC_MAPK1 (PDB:2ERKA, orange) shown in both figures. It is likely the AF2 structures are correct substrate-binding forms of MAPK3 and MAPK7. For STE_MAP3K5, the two dominant PDB conformations (conf1 and conf2) are not typical of substrate-binding structures of STE kinases in the PDB. They also differ substantially from each other in the outer portions of the activation loop. The AF2 models (magenta) are closer to substrate-binding structures of STE kinases in Table 1 (PDB:2q0nA, 4zy4A). For STE_MAP3K14, the AF2 structures are quite different than either of the dominant PDB conformations for reasons that are unknown. TKL_BRAF is not that close to other kinases in the PDB other than RAF1, and the AF2 models are quite different than the benchmark one active structure with a complete activation loop in the PDB (4MNEB). OTHER_BUB1 is also distantly related to all other kinases; in the PDB there are two conformations, one of which has a large bulge blocking the substrate binding site (orange). Most of these structures are not phosphorylated on Ser969, although one of them is (shown in the figure). The other PDB structures (e.g., PDB:4QPM, blue) are more likely to be substrate-capable and most of these structures are phosphorylated on Ser969. The ActiveAF2-template-based model is quite different from both PDB conformations, while the PDB70 and no-template AF2 models (magenta) are much closer to the Active PDB structure (4QPM) and are most likely correct, even though they do not have pLDDT values as high as the ActiveAF2-template structures (green).

**Supplementary Table 1.**
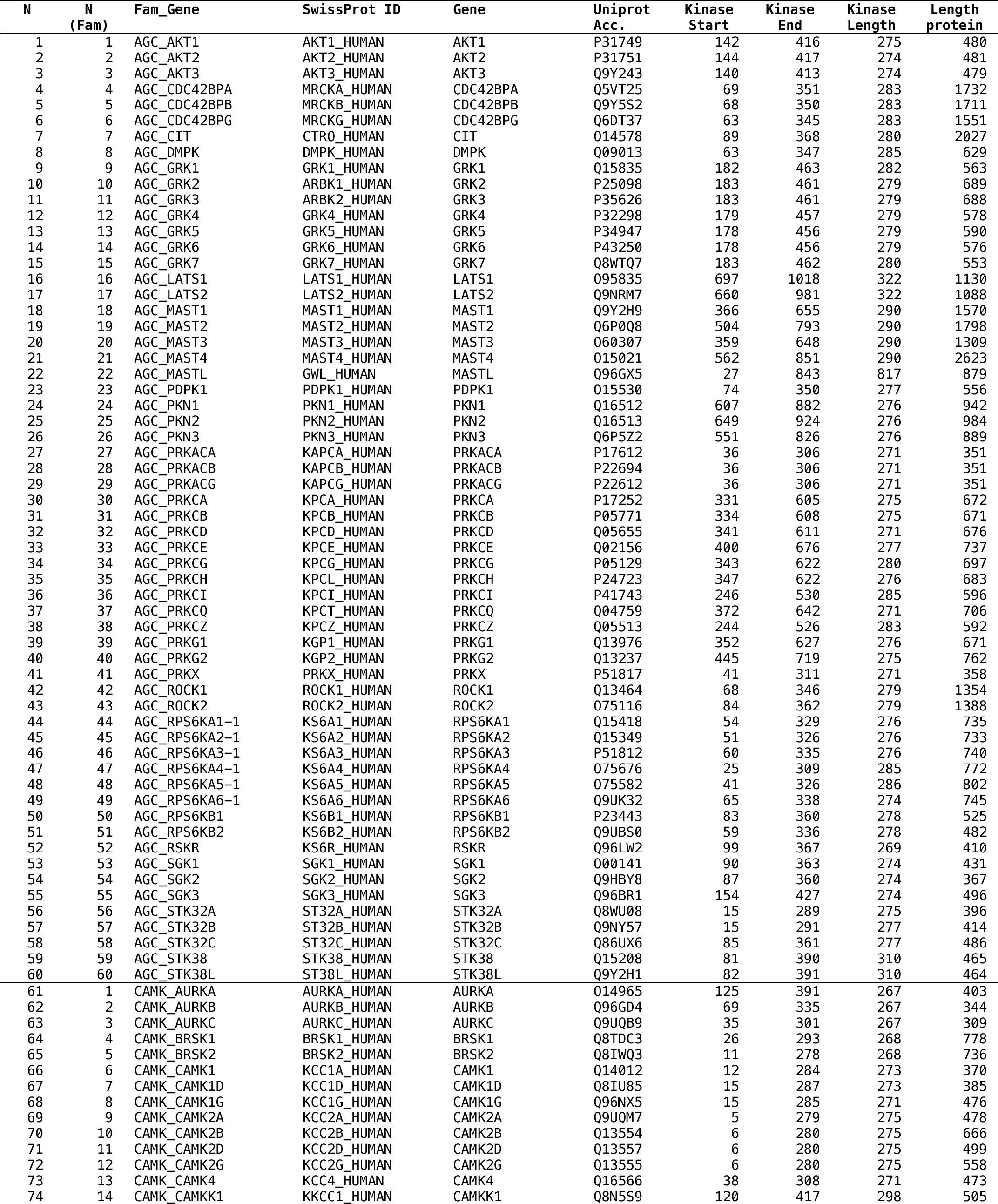
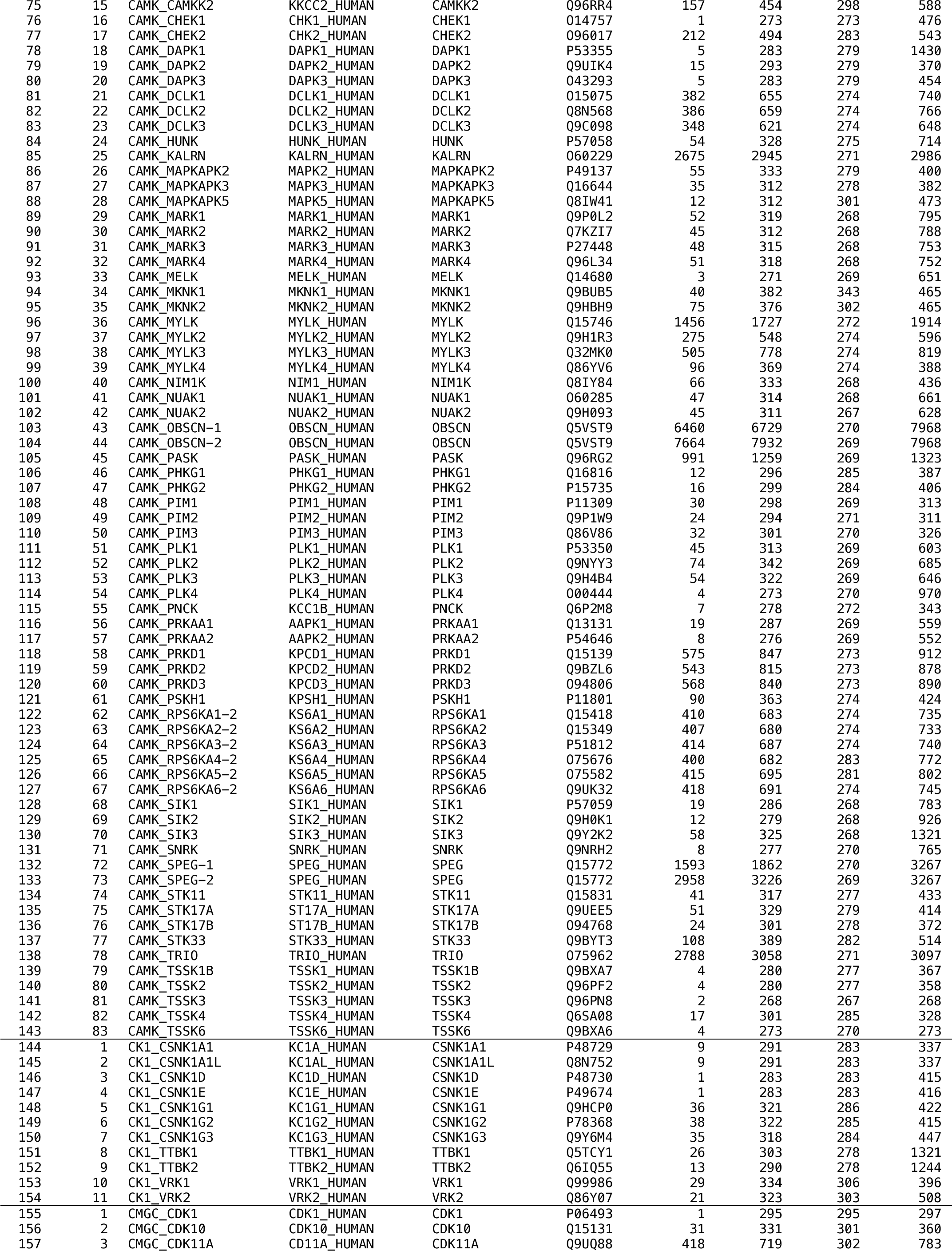
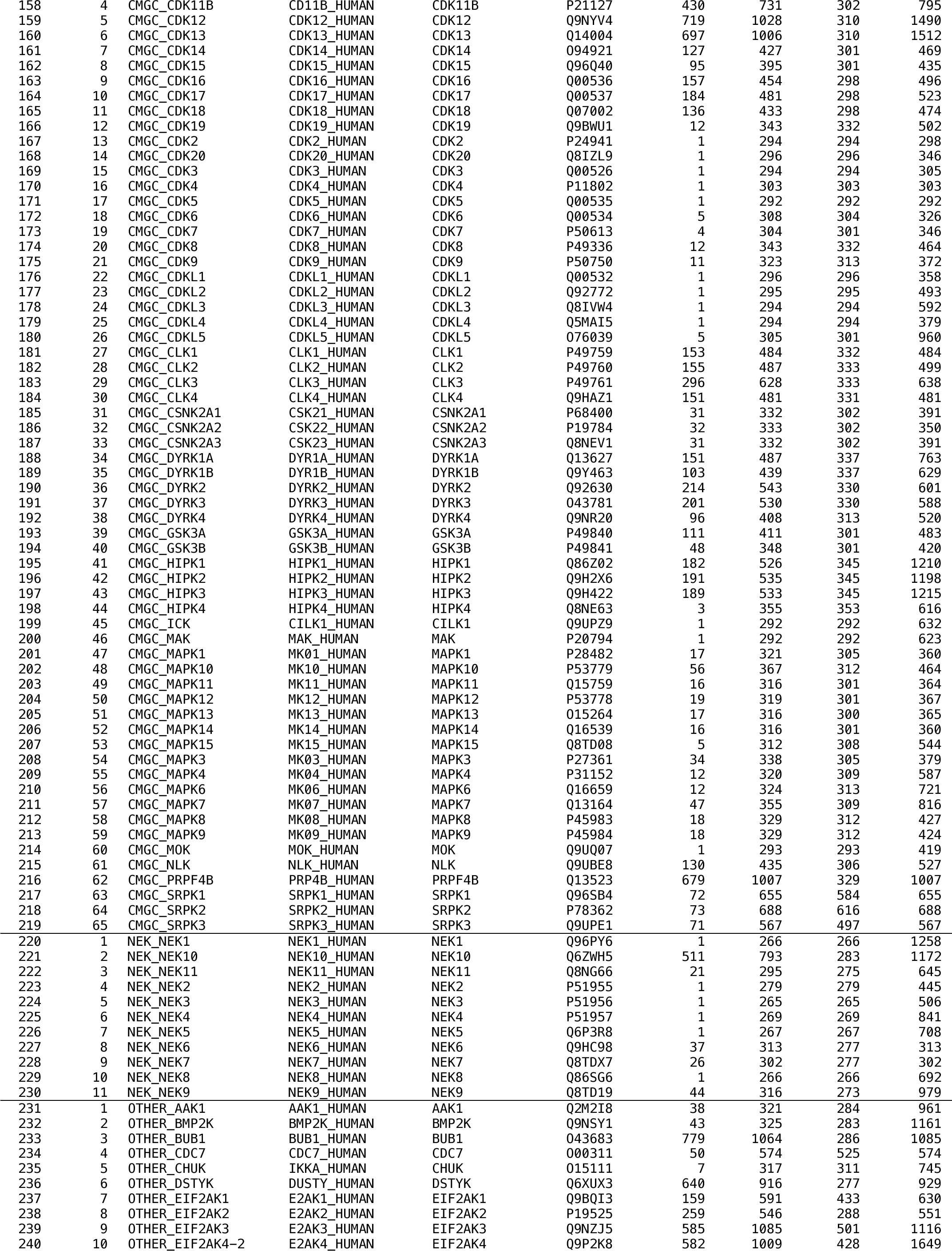
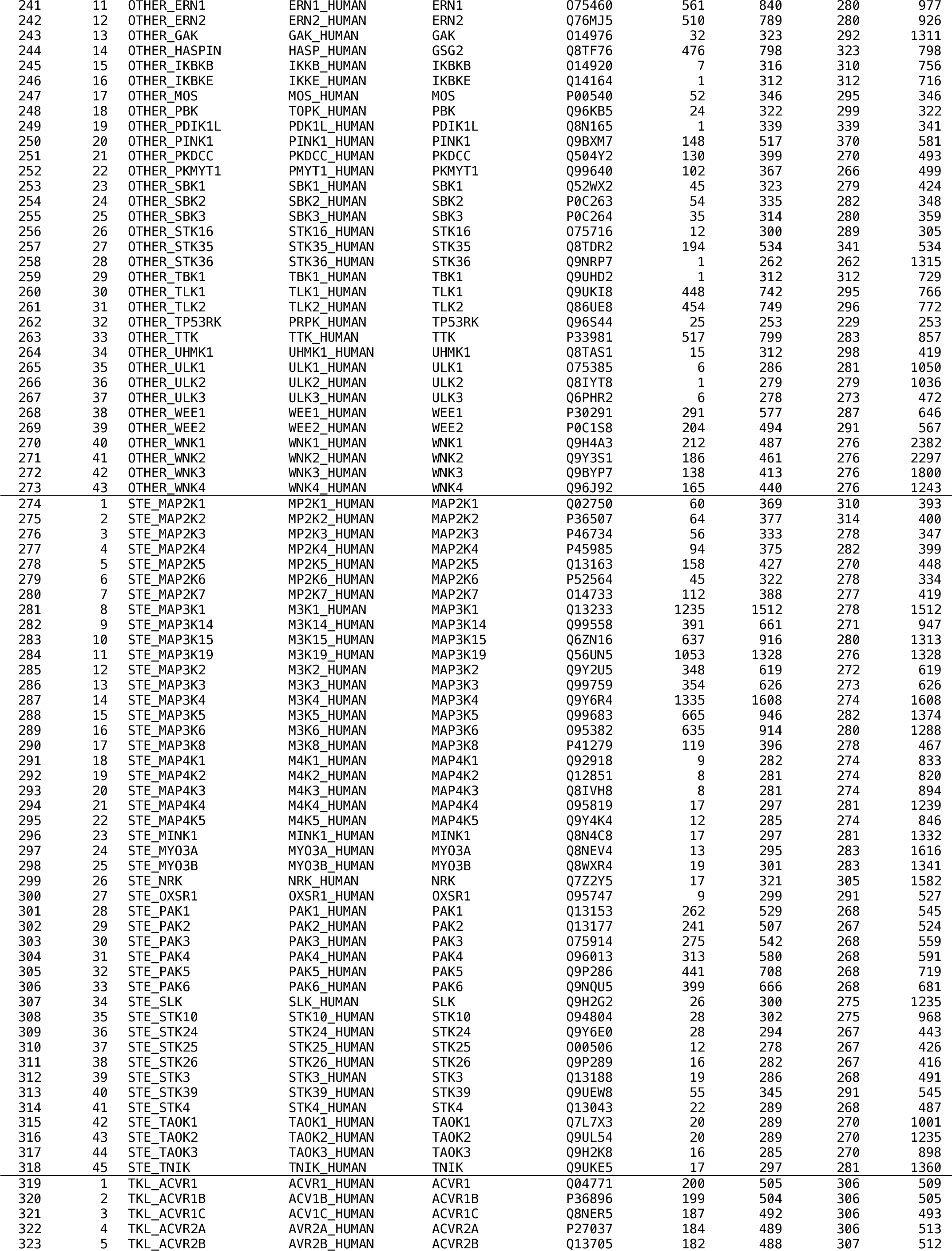
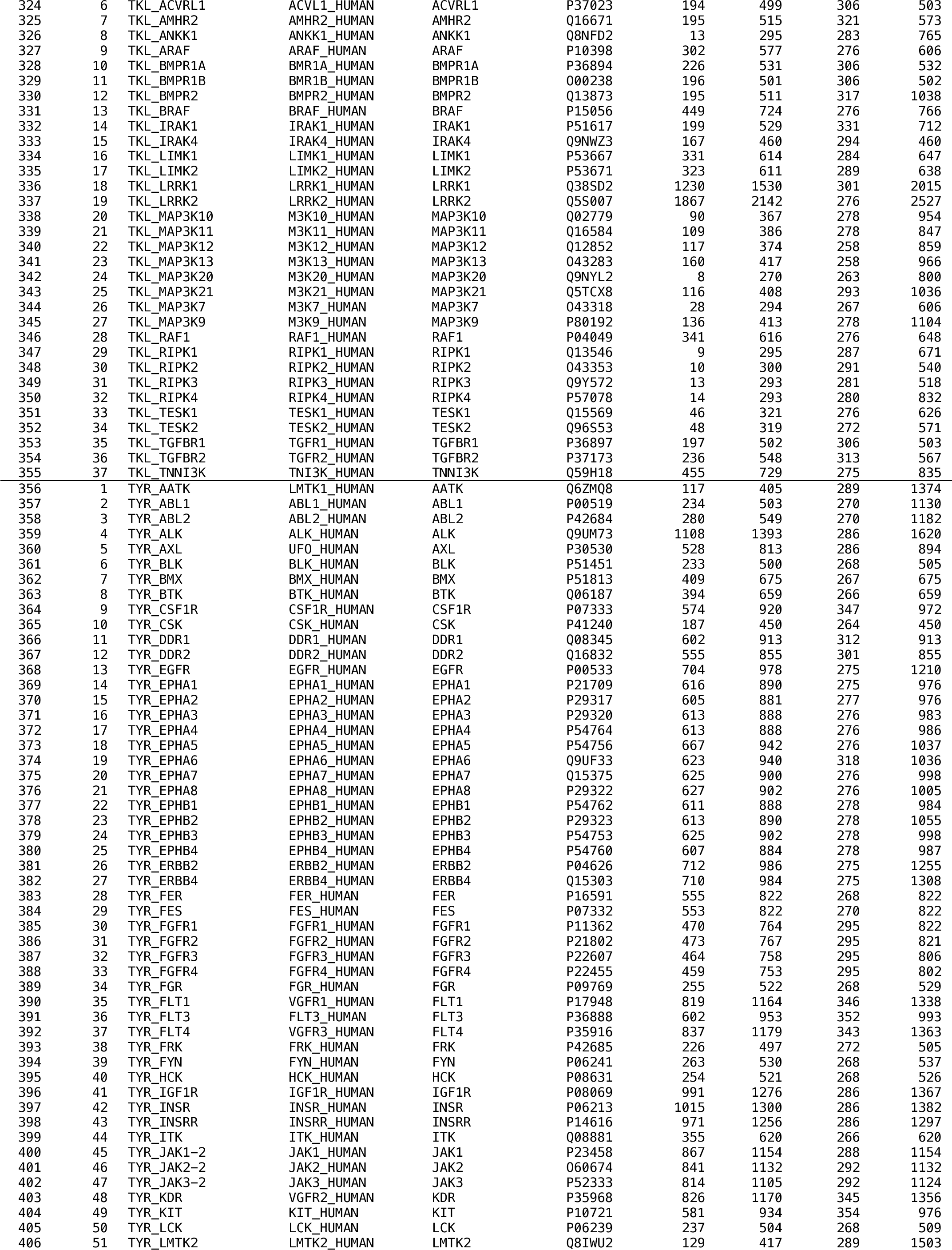
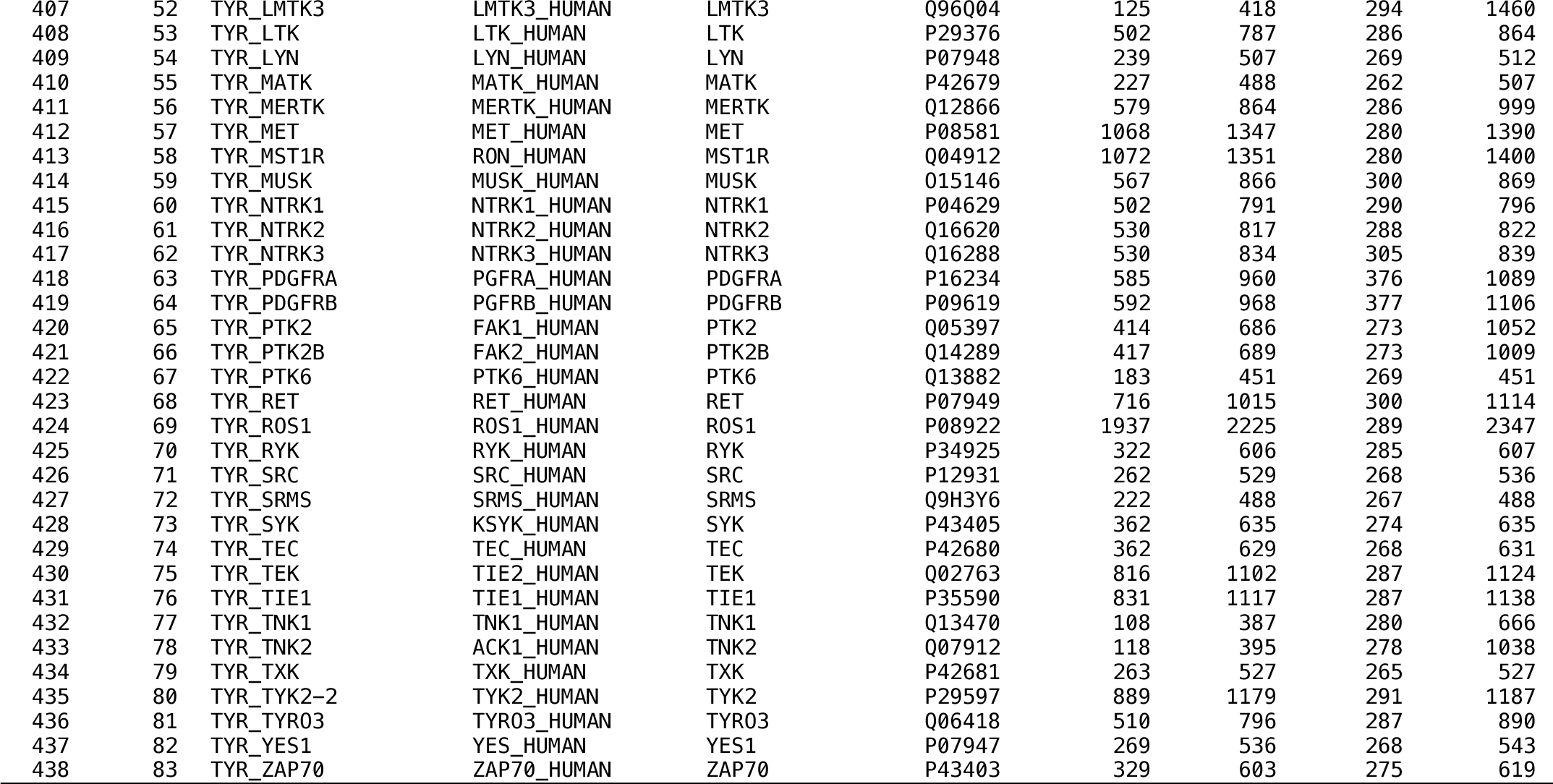
Catalytic kinase domains in the human proteome. The sequence constructs for kinase models produced in this study are given for each human catalytic protein kinase.

**Supplementary Table 2.**
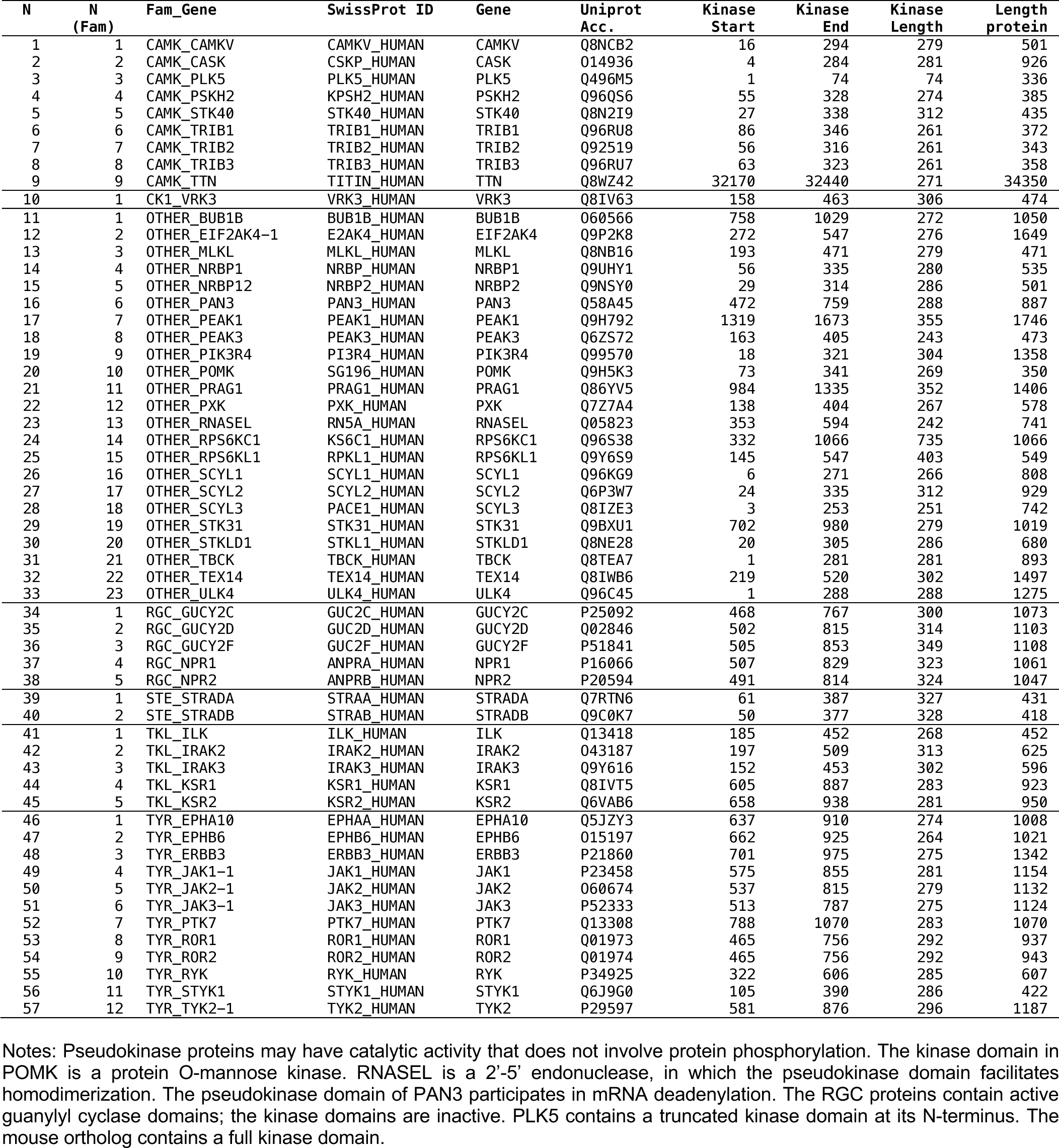
Pseudokinase domains in the human proteome.

